# Natural variation modifies centromere-proximal meiotic crossover frequency and segregation distortion in *Arabidopsis thaliana*

**DOI:** 10.1101/2025.01.04.631303

**Authors:** Nicola Gorringe, Stephanie Topp, Robin Burns, Sota Yamaguchi, Fernando A. Rabanal, Joiselle B. Fernandes, Detlef Weigel, Tetsuji Kakutani, Matthew Naish, Ian R. Henderson

## Abstract

**Background:** Centromeres mediate chromosome segregation during cell division. In plants, centromeres are loaded with CENH3-variant nucleosomes, which direct kinetochore formation and spindle microtubule interaction. Plant centromeres are frequently composed of megabase-scale satellite repeat arrays, or retrotransposon nests. In monocentric genomes, such as *Arabidopsis thaliana*, extended regions of pericentromeric heterochromatin surround the CENH3-occupied satellite arrays. A zone of suppressed meiotic crossover recombination contains the centromere and extends into the pericentromeres. Here, we explore how Arabidopsis natural genetic variation influences centromere-proximal crossover frequency, and segregation distortion through meiosis, when homologous centromeres are structurally heterozygous.

**Results:** We used fluorescent crossover reporters to survey the effect of natural variation on centromere-proximal recombination in twelve F_1_ hybrids, capturing Arabidopsis Eurasian and relict diversity. The majority of F_1_ hybrids showed either elevated or suppressed centromere-proximal crossovers (49 of 60), relative to inbreds. We relate hybrid crossover frequencies to patterns of centromeric structural variation, and in a subset of accessions, to epigenetic patterns of CENH3 loading and DNA methylation. The fluorescent reporters also allow segregation distortion through meiosis to be compared between inbred and hybrid strains. We observed a minority of hybrids (18 of 60) with distorted segregation through meiosis compared to inbreds, with and without simultaneous change to centromere-proximal crossover frequency.

**Conclusions:** We reveal a complex relationship between centromere structural variation, epigenetic state, crossover recombination, and segregation distortion. We propose a model for how Arabidopsis centromere structural heterozygosity may cause segregation distortion during female meiosis.

## Background

Meiosis is a specialised eukaryotic cell division, where a single round of DNA replication is coupled to two rounds of chromosome segregation, which generates a tetrad of spores, each with half the number of chromosomes [1,2]. During meiosis, interhomolog recombination is initiated during prophase I, via the formation of DNA double-strand breaks (DSBs) along the length of chromosomes, by SPO11 complexes [3]. Meiotic DSB formation takes place in the context of a proteinaceous axis - a multiprotein scaffold that organises chromosomes into arrays of tethered chromatin loops [4]. As recombination proceeds, the axes of homologous chromosomes pair, and are held in close physical proximity via the synaptonemal complex [4]. This bivalent arrangement enables meiotic recombination to complete using the homologous chromosome as a repair template, which can result in reciprocal genetic exchange, termed crossover [1,4]. Formation of interhomolog crossovers creates new combinations of genetic variation along chromosomes [1,2], and is additionally required for balanced homolog segregation at the first meiotic division [5].

The frequency of meiotic crossover varies widely along plant chromosomes, with the presence of narrow 1-2 kilobase (kb) hotspots where recombination is elevated over the genome average [6–10]. Plant crossover hotspots are influenced by both DNA sequence and chromatin, and are often found in proximity to euchromatic gene promoters and terminators [6–10]. In the model plant *Arabidopsis thaliana* (hereafter Arabidopsis), the total number of crossovers can be increased by *trans* regulators, for example via overexpression of the E3 ligase HEI10, or mutation of anti-crossover factors [11,12]. Crossovers are also subject to ‘assurance’ and ‘interference’, which respectively describe that each chromosome pair receives an obligate crossover, and that double crossovers are spaced further apart than expected from uniformly random placement [13]. HEI10 coarsening via diffusion along the synaptonemal complex provides a model to understand crossover obligation and interference [14].

Conversely, eukaryotic genomes contain regions of crossover suppression, including heterochromatin, mating-type loci, sex chromosomes, and the centromeres [15–19]. Crossover suppression can be caused both by the inhibitory effects of heterochromatic marks on recombination, as well as sequence polymorphism (heterozygosity) between homologs [15–18]. For example, acquisition of DNA methylation and H3K9me2 methylation can inhibit meiotic crossovers within Arabidopsis recombination hotspots [20]. Equally, heterozygous structural variation has a potent suppressive effect on crossovers [18], and heterochromatic regions, including the centromeres, are highly structurally polymorphic between Arabidopsis accessions [21]. Observations in budding yeast and mammals indicate that centromere-proximal crossovers can be associated with higher rates of gamete aneuploidy [22–24], providing a functional reason to suppress centromere-proximal crossovers. In addition, suppression of crossovers in heterochromatic regions may reduce the chance of illegitimate events, such as unequal crossover, when recombining repetitive templates [25].

Centromeres are defined by the loading of CENH3/CENP-A variant nucleosomes, which direct kinetochore formation and microtubule binding [26,27]. In Arabidopsis, the centromeres consist of megabase arrays of *CEN178* satellite repeats, which are densely DNA methylated [21,28]. The *CEN178* satellite arrays are highly structurally polymorphic between Arabidopsis accessions [21], reflecting the centromere paradox, where centromere sequences rapidly diversify, within and between species, despite playing a conserved and essential role during chromosome segregation [29]. The DNA recombination and repair mechanisms that generate centromere satellite diversity are not well understood but may include somatic and meiotic processes [30,31]. Meiotic recombination maps generated from Arabidopsis Col-0/Ler-0 hybrids showed that the centromeres are contained in megabase-scale non-recombining zones, with proximal crossovers occurring in gene islands embedded in the pericentromeric heterochromatin [17].

An evolutionary explanation for rapid centromere evolution is provided by the drive hypothesis [32–34]. In mammals and plants, female meiosis typically produces a single megaspore/egg that participates in fertilization, while the remaining three sister gametes/spores perish [32–34]. Several genetic systems exist where centromere variants are able to bias their segregation into the surviving female gamete, which causes segregation distortion and non-Mendelian transmission through meiosis [32,35–39]. However, the extent to which intra-species centromere polymorphism causes variation in proximal crossover frequency, and the extent to which this co-occurs with segregation distortion, is not well explored.

Arabidopsis is a powerful model for studying centromere-proximal crossover recombination and segregation distortion, as extensive natural genetic variation in centromere satellite array structures exists, with multiple accessions fully assembled at the sequence level [21,40–42]. Furthermore, fluorescent reporter systems are available that allow sensitive quantification of crossover frequency, and segregation distortion, across the centromeres [17,43,44]. In this study, we generate twelve Arabidopsis hybrids using diverse accessions, crossed to a common background (Col0) that carries centromere-proximal fluorescent transgenic reporters, to measure meiotic crossover frequency and segregation distortion. Compared to inbred strains, the majority of hybrids showed either significantly higher or lower centromere-proximal crossover frequency. We relate these crossover data to centromere structural sequence polymorphism between the recombining chromosomes, and where available, to epigenetic patterns of CENH3 loading and DNA methylation. In a minority of hybrids, we also detected significant segregation distortion of centromere haplotypes through meiosis compared to inbreds, which occurred with or without associated changes to centromere-proximal crossover frequency. We propose a model for how centromere structural polymorphism may cause segregation distortion in Arabidopsis female meiosis.

## Results

### Quantifying centromere-proximal meiotic crossover frequency and segregation distortion using fluorescent reporters in Arabidopsis

In Arabidopsis, segregation of linked, hemizygous T-DNAs that express different colours of fluorescent protein in the pollen or seed can be used to quantify meiotic crossover frequency between the transgene insertions [44–47]. Based on the locations of the Arabidopsis *CEN178* satellite arrays in the Col-CEN genome assembly [28], we selected pairs of FTL T-DNAs that were generated in the Col-0 reference accession [44,46], which define physical intervals that span each of the five centromeres (**Fig. 1A, Fig. S1**, and **Table S1**). In the Col-CEN assembly, the FTL intervals range in size from 4.363 to 7.760 megabases (Mb), and in addition to the *CEN178* satellite arrays, they contain varying amounts of pericentromeric heterochromatin, rDNA copies, genes, and transposable elements (**Fig. 1A, Fig. S1**, and **Table S1**) [28]. When FTL T-DNAs are linked and hemizygous, and the plant is self-fertilized, the number of seeds with each fluorescent phenotype can be used to quantify meiotic crossover frequency (**Fig. 1A and C-D**) [44–47]. When FTL hemizygous plants are self-fertilized, both male and female meiotic crossover are measured at once. In Col-0 inbreds, the centromere-spanning FTL intervals have recombination rates in the range 0.86-2.00 cM/Mb, which are lower than the chromosome arm average of 3.80 cM/Mb measured from Col-0/Ler-0 hybrids [17] (**Table S1**), consistent with centromere-proximal suppression of crossover frequency.

**Figure 1.**
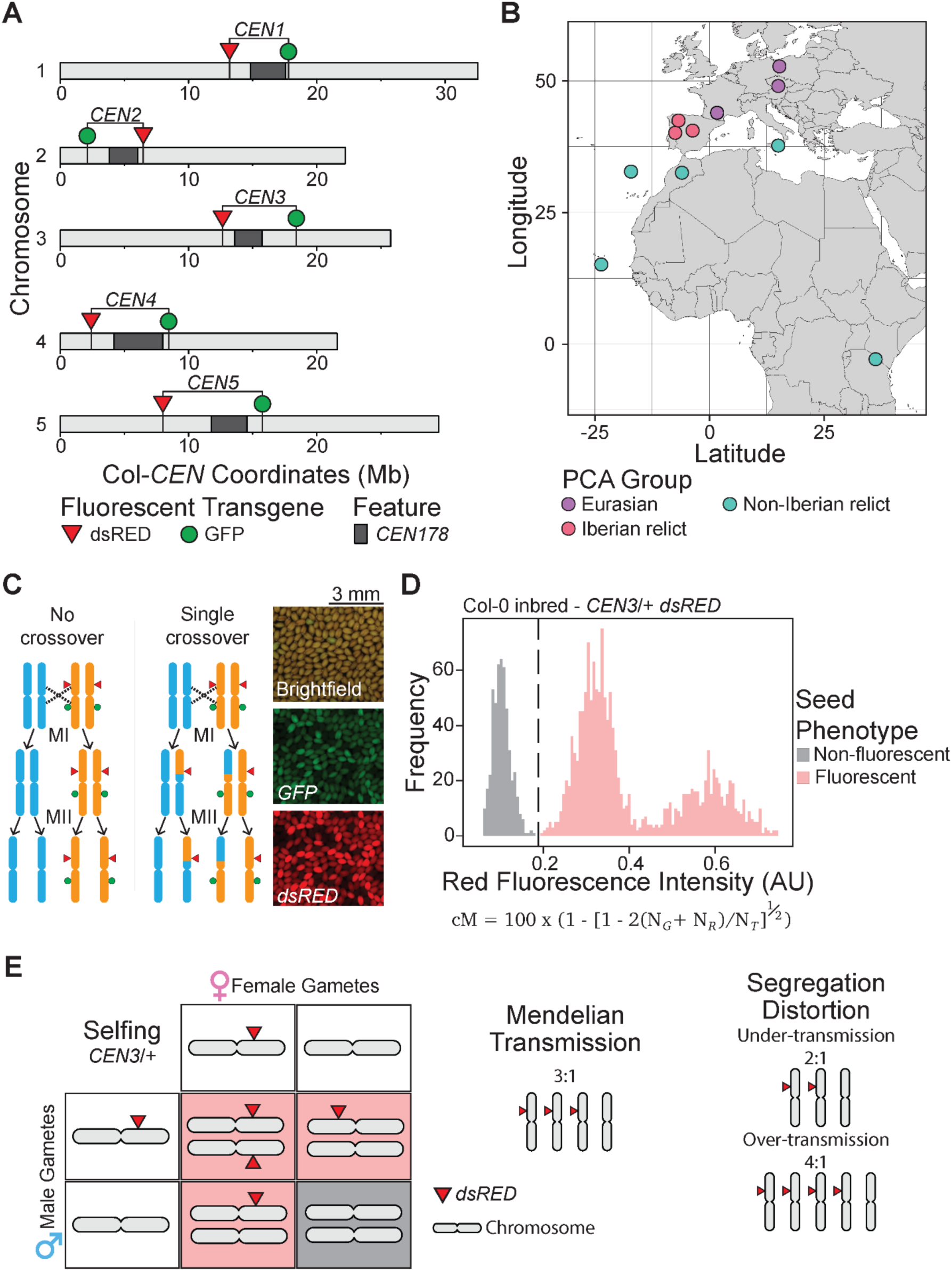
Quantifying centromere-proximal crossover frequency and segregation distortion using fluorescent reporters in Arabidopsis hybrids. **A.** Chromosome ideograms showing the location of FTL T-DNAs used in this study, and the *CEN178* satellite repeat arrays (dark grey), against the Col-CEN genome assembly [28]. Red and green triangles indicate the location of FTL T-DNAs expressing RFP and GFP, respectively [44,46]. **B.** Geographic map showing the origin of the Arabidopsis accessions studied. Accession locations are colour-coded by PCA chromosome arm similarity group [21]; Eurasian (purple), Iberian relict (pink), and Non-Iberian relict (turquoise). **C.** Diagram showing genetic segregation of a hemizygous FTL chromosome through meiosis I (MI) and II (MII). Homologous chromosomes are shown in blue and orange, with the orange genotype representing Col-0 FTL chromosomes, and FTL transgenes shown by red and green triangles. Two segregation scenarios are shown, either with or without a single crossover within the FTL interval. Alongside are representative micrographs of *FTL/+* hemizygous seed imaged in brightfield, or using red or green fluorescence. Scale bar=3 mm. **D.** A representative histogram of red fluorescence (RFP) of seed from a self-fertilized *CEN3/+* hemizygous plant. The dotted line indicates the threshold that divides non-fluorescent (light grey) and fluorescent (red) seed. To calculate crossover frequency (centiMorgan, cM) within the FTL genetic interval, the formula shown is applied to the counts of green (*N_G_*) and red (*N_R_*) fluorescent seed, and the total seed (*N_T_*). **E.** Punnett square showing the expected male and female gametes produced by an FTL hemizygote, and the expected Mendelian inheritance ratio of three fluorescent to one non-fluorescent progeny, following self-fertilization. On the right, is a diagram indicating that higher or lower ratios of fluorescent to non-fluorescent seed can indicate segregation distortion.

The FTL T-DNAs can also be used to quantify patterns of inheritance between generations and segregation distortion. When an FTL hemizygote undergoes meiosis, the Mendelian expectation is that half of the gametes will carry a T-DNA, and half will not (**Fig. 1C** and **1E**). As our F_2_ seed samples represent self-fertilization of male and female meiotic products, the Mendelian expectation is that two thirds of the seed should show fluorescence for either T-DNA; i.e. a third of the seed should be red-fluorescent, and a third should be green-fluorescent (**Fig. 1E**). In the case of segregation distortion, we would expect to observe either an excess of seed with red and green fluorescence if the Col-0 centromere is driving, or fewer coloured seed if the other accession centromere is driving, relative to homozygous inbred backgrounds (**Fig. 1E**). As the centromere is in linkage with the markers, we would expect correlated inheritance behaviour by both red and green FTL markers when centromere drive occurs. In this way, we sought to use the FTL intervals to simultaneously quantify meiotic crossover frequency, and segregation distortion, in centromere-proximal intervals of Arabidopsis hybrids, compared to Col-0 inbred lines.

To investigate the effect of natural variation on FTL crossover frequency in proximity to the centromeres, we selected a diverse panel of twelve accessions for analysis (**Fig. 1B** and **Tables S1** and **S2**). The Arabidopsis accessions selected for study capture diversity from Eurasian (ANGE-B-2, ANGE-B-10, Ler-0, Col-0 and Jm-0), Iberian relict (IP-Cat-0, IP-Med-0 and IP-Alo-19), and non-Iberian relict (Etna-2, Tanz-1, Elk-3, Rabacal-1 and Cvi-0) genome similarity groups (**Fig. 1B** and **Table S2**) [21,28]. The genome assemblies were previously published [21], apart from Jm-0 and Elk-3. Jm-0 was HiFi sequenced to 111× coverage, and Elk-3 was Oxford Nanopore (ONT) sequenced to 1,151× coverage with a subset of >50 kb reads used for assembly (248× coverage). The Jm-0 and Elk-3 genomes were assembled using hifiasm [48]. For Elk-3 all chromosomes were assembled as single contigs, while for Jm-0 nuclear chromosomes there were nine gaps in total. Both assemblies show even long-read alignment coverage across the assembled centromere sequences, and few loci with alternate base signals. For each genome assembly, we used the TRASH algorithm to identify the position and size of the centromeric *CEN178* arrays (**Table S1**) [49].

In our experiments, homozygous FTL lines in the Col-0 accession were crossed with the twelve inbred accessions to generate replicate F_1_ hybrids. The F_1_ hybrids were allowed to self-fertilise and their seed analysed using fluorescence microscopy to quantify both crossover frequency, and segregation distortion (**Fig. 1C-1E** and **Fig. S2**). We first consider each centromere individually, and then together, and relate their genetic and epigenetic organisation to variation in centromere-proximal meiotic crossover frequency and segregation distortion.

### Meiotic crossover frequency and segregation distortion within *CENTROMERE1*

We used the *CTL1.48* interval traffic line [44,46], which consists of a red-fluorescent protein encoding T-DNA *CR1111* located at 13,216,744 base pairs in the Col-CEN genome assembly, and a green-fluorescent protein encoding T-DNA *CG30* located at base pair 17,784,719 (**Fig. 1A, 2A, Fig. S1**, and **Table 1**). For the purposes of this study, we renamed *CTL1.48* as *CENTROMERE1* (*CEN1*), which is 4.567 Mb in the Col-CEN assembly. For each genotype, we quantified *CEN1* crossover frequency (centiMorgans, cM) from multiple replicate individuals, using the counts of red alone (*N_R_*) and green alone (*N_G_*) seeds, compared to the total number (**Fig. 1D** and **Table S3**) [44–47]. To test for significant differences in crossover frequency between the Col-0 inbred controls and each F_1_ hybrid, the *N_R_* and *N_G_* counts were summed across replicates for each genotype and combined with total counts to construct 2×2 contingency tables and Chi-square tests performed.

**Figure 2.**
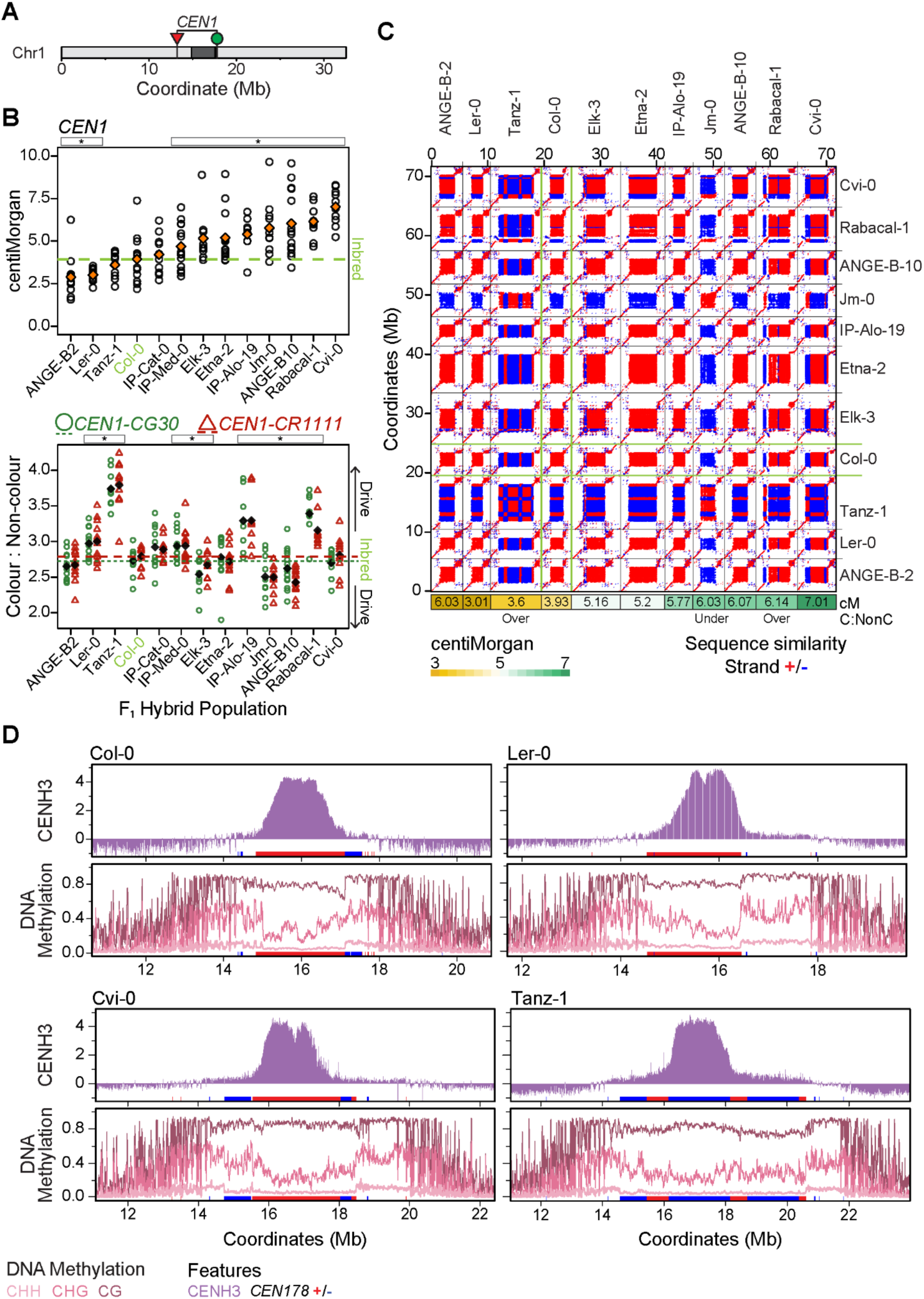
Meiotic crossover frequency, segregation distortion and genetic and epigenetic structure within *CENTROMERE1.* **A.** Chromosome ideogram showing the location of FTL T-DNAs used in this study, together with the position of the *CEN178* satellite repeat arrays (dark grey), against the Col-CEN genome assembly [28]. Red and green triangles indicate the location of FTL T-DNAs expressing RFP and GFP in seed from the *napA* promoter, respectively [44,46]. **B.** Crossover frequency (centiMorgan) values of individual replicates of F_1_ hybrids between the accession listed on the x-axis, and the *CEN1* FTL reporters in a Col-0 background. Mean values are shown in orange diamonds. The inbred Col-0/Col-0-FTL control is highlighted in green, via the horizontal dashed line. Stars at the top of the plot indicate hybrids with significantly different crossover frequencies to the inbred strain based on Chi-square tests. Beneath, the same data are analysed for the ratios of green (circles, short dashed line) and red fluorescent (triangles, long dashed line) seed to non-fluorescent seed, and presented in the same way. **C.** Sequence similarity dot plots comparing the *CEN178* arrays from a subset of the accessions analyzed in B. Red and blue shading indicate sequence similarity using a 161 base pair window. Beneath the dotplot, the cM value for the corresponding *CEN1* hybrid from B. is shown, which is proportionally shaded. Additionally, beneath the dot plot are text indicators showing which accessions showed segregation distortion in B. **D.** For the Col-0 and Ler-0 Eurasian, and Cvi-0 and Tanz-1 relict accessions, we show the CENH3 ChIP-seq enrichment (upper, purple), and DNA methylation (lower) in CG (dark pink), CHG (pink) and CHH (light pink) contexts in the *CEN1* region. The position of *CEN178* arrays are indicated by ticks on the x-axis on the forward (red) or reverse (blue) strand.

Compared to Col-0 inbreds, we observed that ANGE-B-2 and Ler-0 F_1_ hybrids showed significantly lower crossover frequency, whereas eight hybrids were significantly higher (**Fig. 2B**, **Table S1**, and **Table S3**). The two highest-recombining hybrids, Rabacal-1 and Cvi-0, are diverged relict accessions compared to the Eurasian Col-0 parent, with larger arrays *CEN178* centromere arrays to Col-0 by 8,523 and 7,717 copies, respectively; whereas, the low-recombining ANGE-B-2 and Ler-0 hybrids have *CEN178* arrays more similar in size and structure to Col-0 (**Fig. 2B-2C**, **Table S1**, and **Table S3**). Further, taken together there was no significant correlation between crossover frequency and the size or similarity of the hybrid *CEN178* arrays (**Fig. 2B–2C**, **Fig. S3**, and **Table S1**). For example, the Tanz-1 accession, which has the largest *CEN178* array (34,131 copies), which are arranged in series of inverted blocks, has a crossover frequency not significantly different to Col-0 inbred (**Fig. 2B-2C**, **Fig. S4**, and **Table S1**). Jm-0 was also notable as it bears a *CEN178* array on the opposite strand to Col-0 and yet, has significantly higher crossover frequency (**Fig. 2B-2C**, **Fig. S4**, and **Table S1**). We conclude that although there is substantial variation in *CEN1* centromere-proximal crossover frequency in Arabidopsis hybrids, this is not correlated with structural variation in the *CEN178* arrays.

Using the FTL seed data, we next calculated the ratio of red (*N_R_*) to non-red fluorescent (*N_NR_*), and green (*N_G_*) to non-green fluorescent (*N_NG_*) seed. As expected, we observed a strong correlation between red and green fluorescence ratios within genotypes for all *CEN* intervals (**Fig. 2B**, **Fig. S2**, **Table S1**, and **Table S3**). For each F_1_ hybrid sample we compared the fluorescent and non-fluorescent seed counts to inbreds, using 2×2 contingency tables and performed Chi-square tests to assess significance (**Fig. 2B**). When both red and green inheritance ratios are significantly different to Col-0 inbreds, with the same direction of change, we consider this as evidence of segregation distortion. We observed that five F_1_ hybrids; Ler-0, Tanz-1, IP-Med-0, IP-Alo-19 and Rab-1, showed significant over-transmission of the Col-0 centromere *CEN1* interval, whereas three; Elk-3, Jm-0, and ANGE-B-10, showed significant under-transmission, compared to inbreds (**Fig. 2B-2C** and **Fig. S2**). The direction and strength of distorted inheritance did not correlate with changes to meiotic crossover frequency. For example, the strongest drive against Col-0 occurred in Tanz-1 hybrids, where there was no change to crossover frequency, as well as in the highly recombining IP-Alo-19 and Rab-1 hybrids (**Fig. 2B**, **Table S1**, and **Table S3**). Equally, there was no correlation between the strength or direction of distorted inheritance and how different hybrid *CEN178* arrays were in terms of size and similarity (**Fig. 2B-2C**, **Fig. S3-S4**, and **Table S1**). Together, this reveals distorted inheritance of *CEN1* in Arabidopsis hybrids, compared to inbreds, but which did not correlate with either *CEN178* structural variation, or proximal crossover frequency.

As *CEN1* structural polymorphism was not strongly correlated with variation in crossover frequency, we examined epigenetic organisation. In a subset of accessions; the Eurasian Col-0 and Ler-0, and the relicts Cvi-0 and Tanz-0, we compared ChIP-seq enrichment of the centromeric histone CENH3, together with DNA methylation in CG, CHG and CHH sequence contexts across *CEN1* [21]. As previously reported, CENH3 showed a consistent peak of enrichment ∼1 Mb in width in all accessions, despite significant variation in *CEN178* array size and structure (**Fig. 2D**). In each accession, the centromeres are embedded in highly DNA methylated pericentromeric regions (**Fig. 2D**). The *CEN178* arrays themselves show elevated CG context DNA methylation, but reduced CHG and CHH methylation, relative to the peri-centromeres (**Fig. 2D**). In some cases, such as Tanz-1, CENH3 loading correlates with the underlying DNA sequence, in this case being enriched within a single *CEN178* inversion block (**Fig. 2D**). Therefore, patterns of epigenetic organisation between the recombining centromeres do not clearly explain variation in centromere-proximal crossover frequency, or distorted inheritance.

### Meiotic crossover frequency and segregation distortion within *CENTROMERE2*

For chromosome 2, we used the *CTL2.16* traffic line [44,46], which consists of a red fluorescent protein encoding T-DNA *CR1014* located at 6,472,440 base pairs in the Col-CEN genome assembly, and a green-fluorescent protein encoding T-DNA *CG531* located at coordinate 2,108,979. For the purposes of this study, we renamed *CTL2.16* as *CENTROMERE2* (*CEN2*), which is 4.363 Mb in the Col-CEN assembly (**Fig. 1A**, **3A**, **Fig. S1**, and **Table S1**). In *CEN2*, we observed a single hybrid (Ler-0) that had significantly lower crossover frequency than inbreds, whereas nine were significantly higher (**Fig. 3B**, **Table S1**, and **Table S4**). As for *CEN1*, we did not observe a significant correlation between *CEN178* array size, or structural polymorphism, and *CEN2* crossover frequency in the hybrids (**Fig. 3B-3C**, **Fig. S3**, **S5**, and **Table S1**). In *CEN2* hybrids with the divergent relict Cvi-0, Tanz-1 and IP-Med-0 accessions, which have polymorphic *CEN178* arrays, we observed significantly elevated hybrid crossovers, compared to inbreds, indicating again that centromeric genetic divergence is not a barrier to elevated recombination (**Fig. 3B-3C**, **Fig. S5**, and **Table S1**).

**Figure 3.**
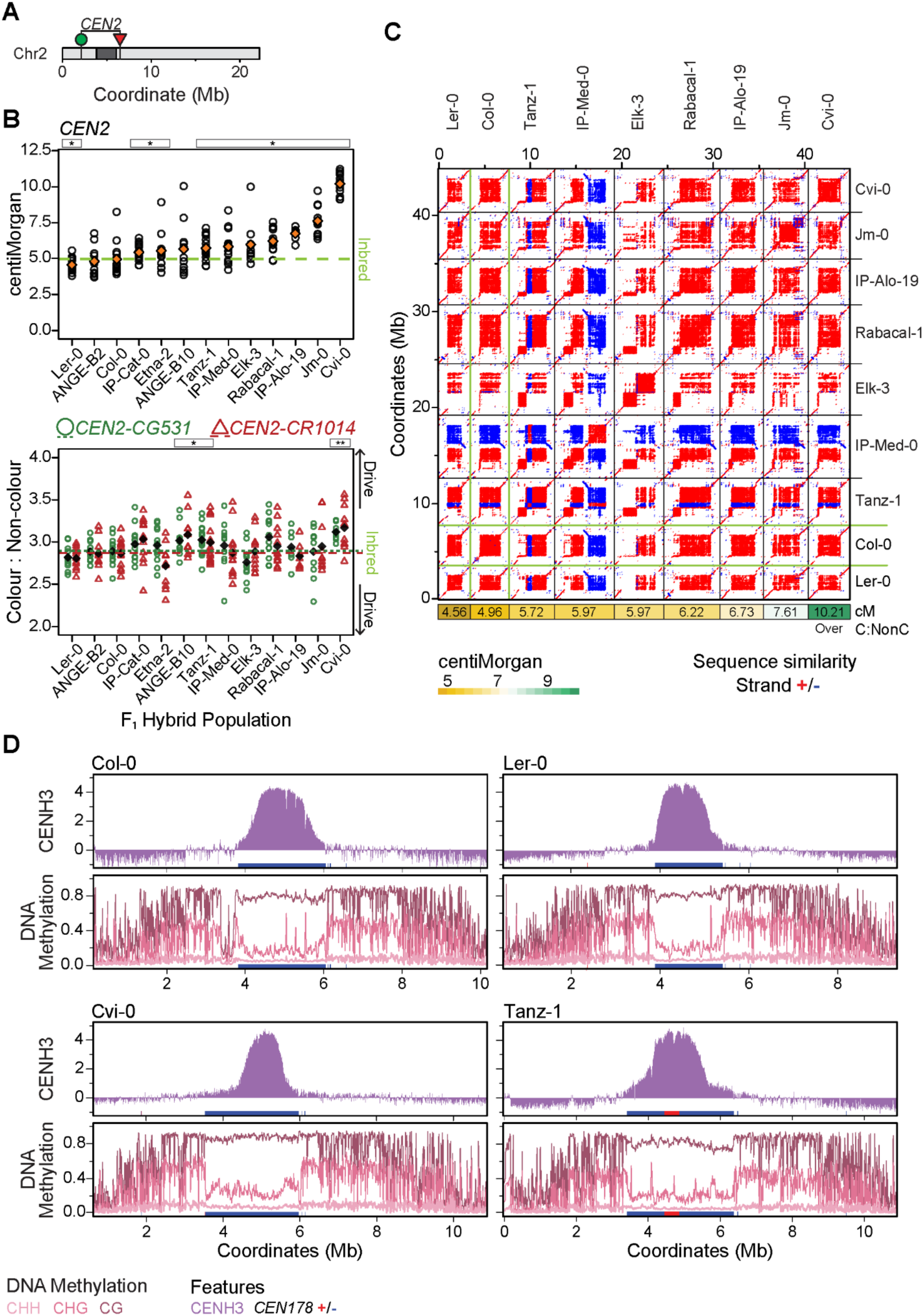
Meiotic crossover frequency, segregation distortion and genetic and epigenetic structure within *CENTROMERE2.* **A.** Chromosome ideogram showing the location of FTL T-DNAs used in this study, together with the position of the *CEN178* satellite repeat arrays (dark grey), against the Col-CEN genome assembly [28]. Red and green triangles indicate the location of FTL T-DNAs expressing RFP and GFP in seed from the *napA* promoter, respectively [44,46]. **B.** Crossover frequency (centiMorgan) values of individual replicates of F_1_ hybrids between the accession listed on the x-axis, and the *CEN2* FTL reporters in a Col-0 background. Mean values are shown in orange diamonds. The inbred Col-0/Col-0-FTL control is highlighted in green, via the horizontal dashed line. Stars at the top of the plot indicate hybrids with significantly different crossover frequencies to the inbred strain based on Chi-square tests. Beneath, the same data are analysed for the ratios of green (circles, short dashed line) and red (triangles, long dashed line) fluorescent seed to non-fluorescent seed, and presented in the same way. **C.** Sequence similarity dot plots comparing the *CEN178* arrays from a subset of the accessions analyzed in B. Red and blue shading indicate sequence similarity using a 161 base pair window. Beneath the dotplot, the cM value for the corresponding *CEN2* hybrid from B. is shown, which is proportionally shaded. Additionally, beneath the dot plot are text indicators showing which accessions showed segregation distortion in B. **D.** For the Col-0 and Ler-0 Eurasian, and Cvi-0 and Tanz-1 relict accessions, we show the CENH3 ChIP-seq enrichment (upper, purple), and DNA methylation (lower) in CG (dark pink), CHG (pink) and CHH (light pink) contexts in the *CEN2* region. The position of *CEN178* arrays are indicated by ticks on the x-axis on the forward (red) or reverse (blue) strand.

Compared to *CEN1*, fewer of the *CEN2* hybrids showed significant segregation distortion, with only ANGE-B-10, Tanz-1 and Cvi-0 hybrids showing over-transmission of the Col-0 centromere (**Fig. 3B**, **Fig. S5**, and **Table S1**). Similar to *CEN1*, the incidence of segregation distortion did not correlate either with the degree of *CEN178* array polymorphism, or the frequency of centromere-proximal crossover frequency (**Fig. 3B-3C**, **Fig. S3**, **S5**, **Table S1**, and **Table S4**). In each of the four highlighted accessions, within *CEN2*, there exists a ∼1 Mb region of CENH3 loading with elevated CG DNA methylation and depleted CHG and CHH methylation in the *CEN178* arrays relative to the peri-centromeres (**Fig. 3D**). Comparison of the epigenetic landscape of the Col-0 centromere, with the driving Cvi-0 and Tanz-1 centromeres, did not reveal an obvious difference (**Fig. 3D**).

### Meiotic crossover frequency and segregation distortion within *CENTROMERE3*

To study centromere 3, we combined the red-encoding FTL T-DNA *CR880*, located at 12,652,136 base pairs in the Col-CEN genome assembly, with a green-encoding T-DNA *CG811* at base pair 18,395,802, and named this interval *CENTROMERE3* (*CEN3*), which is 5.743 Mb (**Fig. 1A**, **4A**, **Fig. S1**, and **Table S1**) [44,46]. In contrast to *CEN1* and *CEN2*, where the majority of hybrids showed significantly higher crossover frequency compared to inbreds, in *CEN3*, six hybrids were lower, and two were higher (**Fig. 4B**, **Table S1**, and **Table S5**). Interestingly, the Cvi-0 and IP-Alo-19 relict accessions again showed elevated crossover frequency, despite structural polymorphism in the *CEN178* arrays, compared to Col-0 (**Fig. 4B-4C**, **Fig. S3**, **S6**, and **Table S1**). In Col-0, the *CEN3 CEN178* arrays have an inverted array structure [28], that is shared to varying degrees with Elk-3, Ler-0, Rab-1, Cvi-0, and IP-Alo-19, and which together include the highest and lowest recombining hybrids (**Fig. 4B-4C**, **Fig. S6** and **Table S1**). Three relict hybrids; Elk-3, Tanz-1, and Rab-1, showed significant over-transmission of the Col-0 centromere, whereas the Eurasian Jm-0 hybrid showed under-transmission (**Fig. 4B**, **Fig. S6** and **Table S1**). With respect to CENH3 loading, in Cvi-0, Ler-0 and Tanz-1 ChIP-seq enrichment was observed within specific *CEN178* sub-arrays, whereas in Col-0 loading was found to span both inverted arrays (**Fig. 4D**). As for *CEN1* and *CEN2*, variation in *CEN3* crossover frequency was uncoupled from the incidence of segregation distortion, and was not correlated with the degree of centromeric repeat array variation, or epigenetic features.

**Figure 4.**
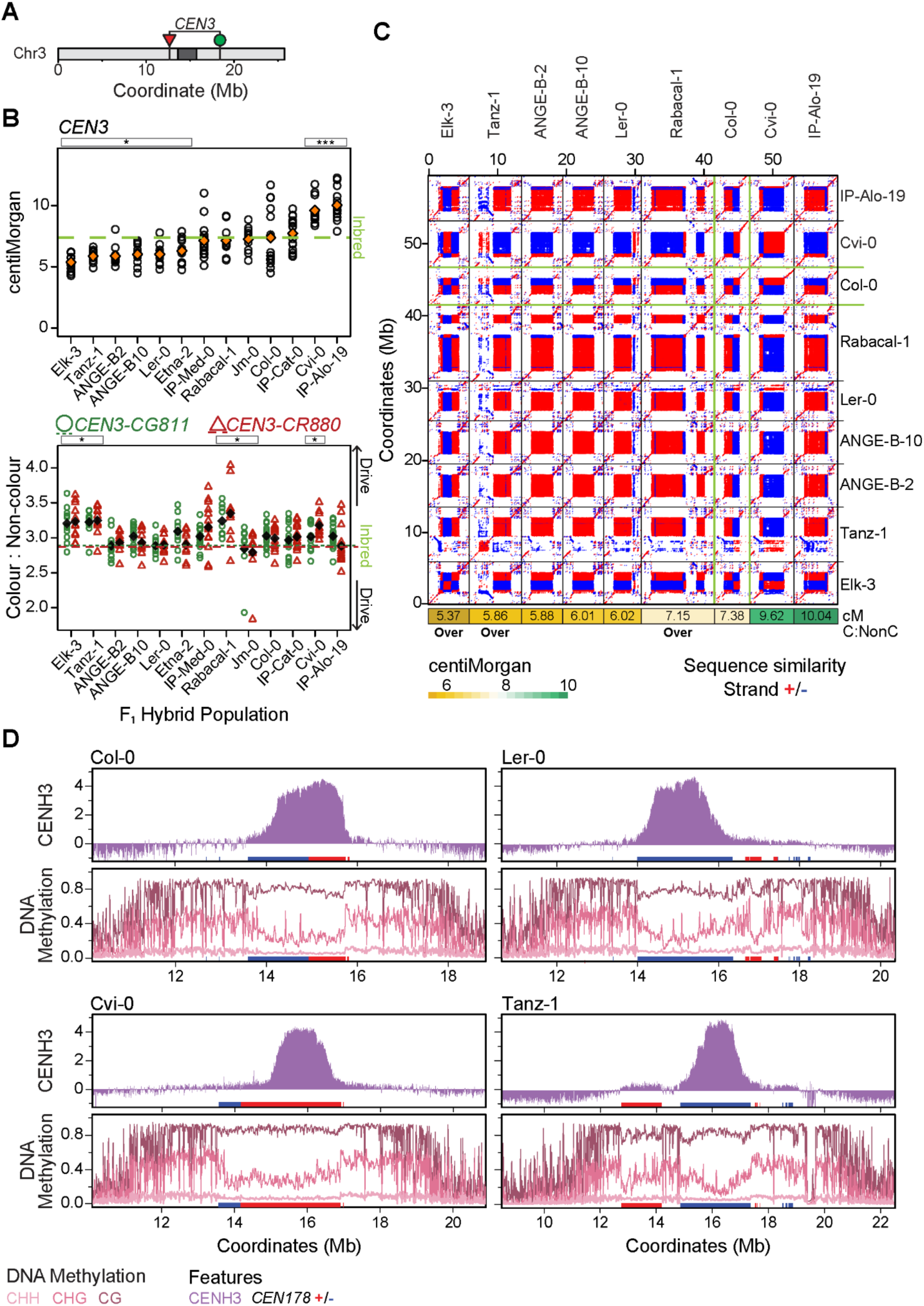
Meiotic crossover frequency, segregation distortion and genetic and epigenetic structure within *CENTROMERE3.* **A.** Chromosome ideogram showing the location of FTL T-DNAs used in this study, together with the position of the *CEN178* satellite repeat arrays (dark grey), against the Col-CEN genome assembly [28]. Red and green triangles indicate the location of FTL T-DNAs expressing RFP and GFP in seed from the *napA* promoter, respectively [44,46]. **B.** Crossover frequency (centiMorgan) values of individual replicates of F_1_ hybrids between the accession listed on the x-axis, and the *CEN3* FTL reporters in a Col-0 background. Mean values are shown in orange diamonds. The inbred Col-0/Col-0-FTL control is highlighted in green, via the horizontal dashed line. Stars at the top of the plot indicate hybrids with significantly different crossover frequencies to the inbred strain based on Chi-square tests. Beneath, the same data are analysed for the ratios of green (circles, short dashed line) and red (triangles, long dashed line) fluorescent seed to non-fluorescent seed, and presented in the same way. **C.** Sequence similarity dot plots comparing the *CEN178* arrays from a subset of the accessions analyzed in B. Red and blue shading indicate sequence similarity using a 161 base pair window. Beneath the dotplot, the cM value for the corresponding *CEN3* hybrid from B. is shown, which is proportionally shaded. Additionally, beneath the dot plot are text indicators showing which accessions showed segregation distortion in B. **D.** For the Col-0 and Ler-0 Eurasian, and Cvi-0 and Tanz-1 relict accessions, we show the CENH3 ChIP-seq enrichment (upper, purple), and DNA methylation (lower) in CG (dark pink), CHG (pink) and CHH (light pink) contexts in the *CEN3* region. The position of *CEN178* arrays are indicated by ticks on the x-axis on the forward (red) or reverse (blue) strand.

**Figure 5.**
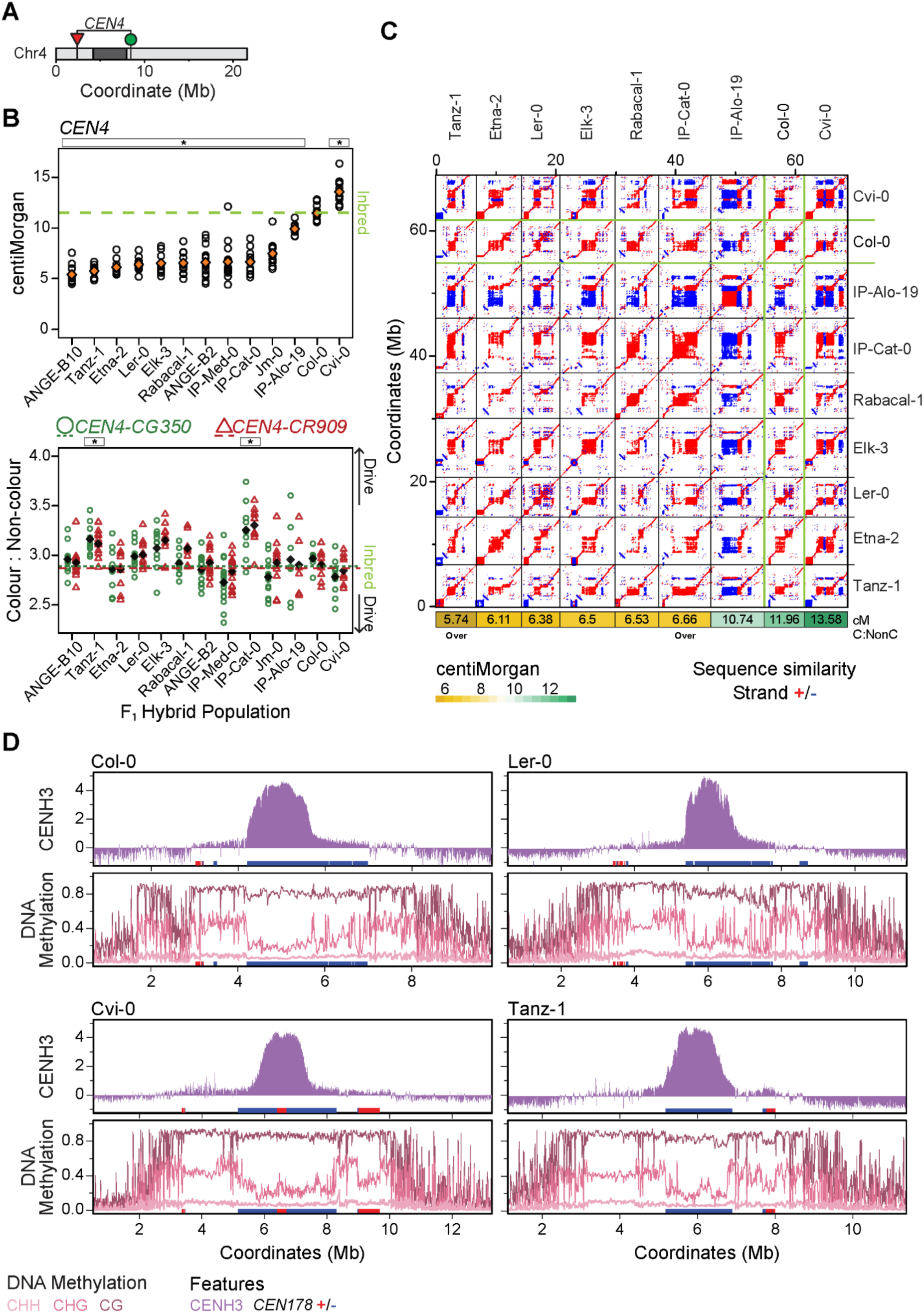
Meiotic crossover frequency, segregation distortion and genetic and epigenetic structure within *CENTROMERE4.* **A.** Chromosome ideogram showing the location of FTL T-DNAs used in this study, together with the position of the *CEN178* satellite repeat arrays (dark grey), against the Col-CEN genome assembly [28]. Red and green triangles indicate the location of FTL T-DNAs expressing RFP and GFP in seed from the *napA* promoter, respectively [44,46]. **B.** Crossover frequency (centiMorgan) values of individual replicates of F_1_ hybrids between the accession listed on the x-axis, and the *CEN4* FTL reporters in a Col-0 background. Mean values are shown in orange diamonds. The inbred Col-0/Col-0-FTL control is highlighted in green, via the horizontal dashed line. Stars at the top of the plot indicate hybrids with significantly different crossover frequencies to the inbred strain based on Chi-square tests. Beneath, the same data are analysed for the ratios of green (circles, short dashed line) and red (triangles, long dashed line) fluorescent seed to non-fluorescent seed, and presented in the same way. **C.** Sequence similarity dot plots comparing the *CEN178* arrays from a subset of the accessions analyzed in B. Red and blue shading indicate sequence similarity using a 161 base pair window. Beneath the dotplot, the cM value for the corresponding *CEN4* hybrid from B. is shown, which is proportionally shaded. Additionally, beneath the dot plot are text indicators showing which accessions showed segregation distortion in B. **D.** For the Col-0 and Ler-0 Eurasian, and Cvi-0 and Tanz-1 relict accessions, we show the CENH3 ChIP-seq enrichment (upper, purple), and DNA methylation (lower) in CG (dark pink), CHG (pink) and CHH (light pink) contexts in the *CEN4* region. The position of *CEN178* arrays are indicated by ticks on the x-axis on the forward (red) or reverse (blue) strand.

**Figure 6.**
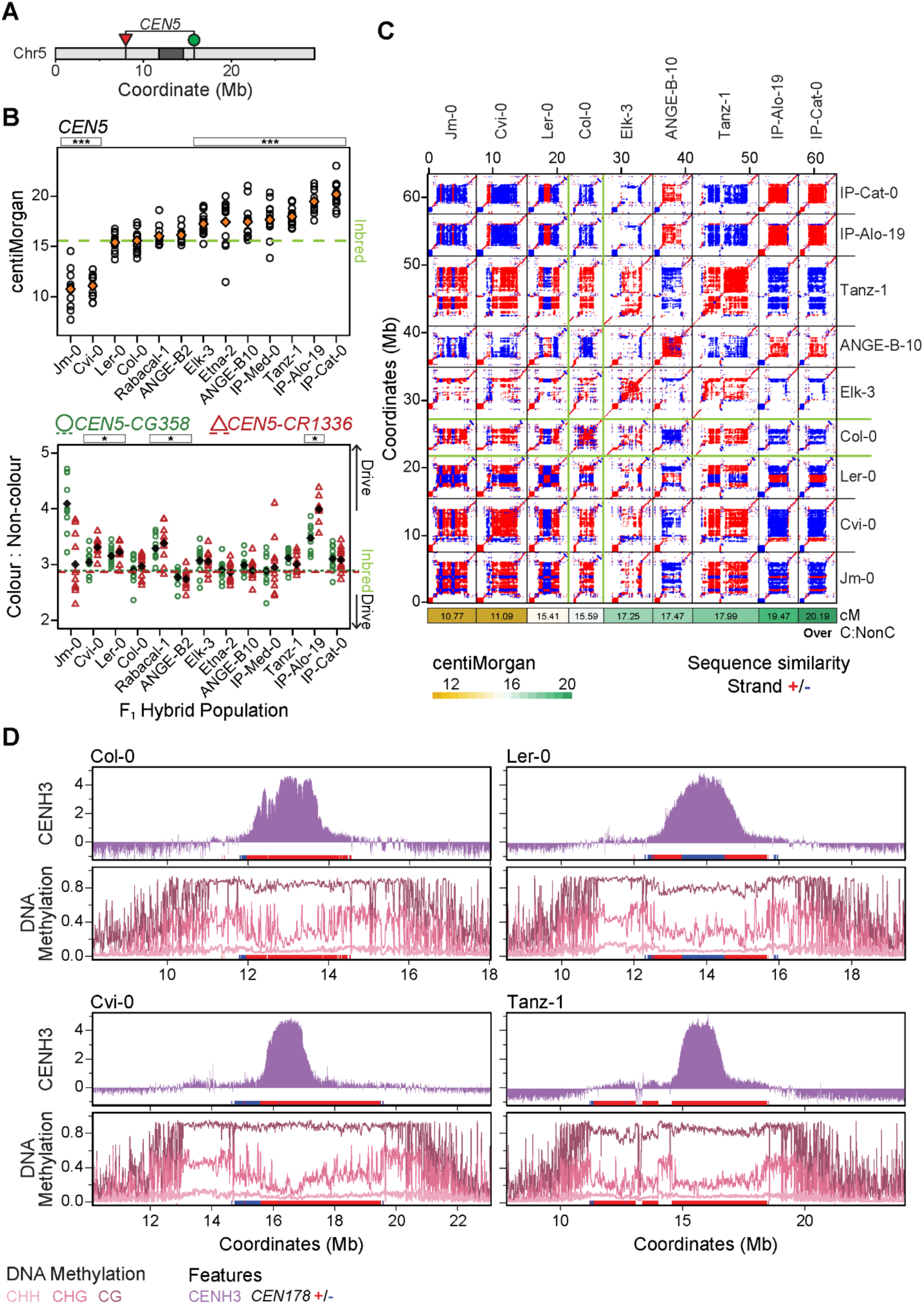
Meiotic crossover frequency, segregation distortion and genetic and epigenetic structure within *CENTROMERE5.* **A.** Chromosome ideogram showing the location of FTL T-DNAs used in this study, together with the position of the *CEN178* satellite repeat arrays (dark grey), against the Col-CEN genome assembly [28]. Red and green triangles indicate the location of FTL T-DNAs expressing RFP and GFP in seed from the *napA* promoter, respectively [44,46]. **B.** Crossover frequency (centiMorgan) values of individual replicates of F_1_ hybrids between the accession listed on the x-axis, and the *CEN5* FTL reporters in a Col-0 background. Mean values are shown in orange diamonds. The inbred Col-0/Col-0-FTL control is highlighted in green, via the horizontal dashed line. Stars at the top of the plot indicate hybrids with significantly different crossover frequencies to the inbred strain based on Chi-square tests. Beneath, the same data are analysed for the ratios of green (circles, short dashed line) and red (triangles, long dashed line) fluorescent seed to non-fluorescent seed, and presented in the same way. **C.** Sequence similarity dot plots comparing the *CEN178* arrays from a subset of the accessions analyzed in B. Red and blue shading indicate sequence similarity using a 161 base pair window. Beneath the dotplot, the cM value for the corresponding *CEN5* hybrid from B. is shown, which is proportionally shaded. Additionally, beneath the dot plot are text indicators showing which accessions showed segregation distortion in B. **D.** For the Col-0 and Ler-0 Eurasian, and Cvi-0 and Tanz-1 relict accessions, we show the CENH3 ChIP-seq enrichment (upper, purple), and DNA methylation (lower) in CG (dark pink), CHG (pink) and CHH (light pink) contexts in the *CEN5* region. The position of *CEN178* arrays are indicated by ticks on the x-axis on the forward (red) or reverse (blue) strand.

### Meiotic crossover frequency and segregation distortion within *CENTROMERE4*

For chromosome four, we used the *CTL4.12* interval traffic line [44,46], which consists of a red-encoding T-DNA *CR909* at coordinate 2,437,418 in the Col-CEN genome assembly, and a green-encoding T-DNA *CG350* at coordinate 8,480,638. For the purposes of this study, we renamed *CTL4.12* as *CENTROMERE4* (*CEN4*), which is 6.043 Mb in width (**Fig. 1A**, **5A**, **Fig. S1** and **Table S1**). For *CEN4*, the majority of hybrids (11 of 12) showed significantly reduced crossover frequency compared to inbreds, with the only exception being the Cvi-0 relict hybrid showing significantly higher recombination (**Fig. 5B**, **Table S1**, and **Table S6**). Col-0 is documented to carry a large ‘knob’ inversion on the short arm of chromosome four that spans the *CR909* FTL T-DNA [50,51], which was absent in the twelve other accessions crossed to Col-0 (**Fig. S7**), and likely contributes to suppressed crossover frequency in the *CEN4* hybrids. Two relict accessions, Tanz-1 and IP-Cat-0, showed distorted inheritance in favour of the Col-0 chromosome (**Fig. 5B**, **Table S1**, and **Table S6**). The accessions profiled at the level of chromatin showed the presence of multiple distinct *CEN178* arrays, only one of which was CENH3-occupied in each case (**Fig. 5D**).

### Meiotic crossover frequency and segregation distortion within *CENTROMERE5*

We crossed the red-encoding FTL T-DNA *CR1136* at coordinate 7,998,504 in the Col-CEN genome assembly, and a green-encoding FTL T-DNA *CG358* at coordinate 15,759,366, to generate the *CENTROMERE5* (*CEN5*) interval, which is 7.760 Mb in width (**Fig. 1A**, **6A, Fig. S1**, and **Table S1**) [44,46]. Compared to the other *CEN* FTL intervals, *CEN5* contains a higher proportion of pericentromeric sequence in addition to the *CEN178* satellite arrays (**Fig. 1A**, **S1**, and **Table S1**). In *CEN5*, seven hybrids showed significantly higher crossover frequency compared to inbreds, and two were significantly lower, with the relict hybrids IP-Alo-19 and IP-Cat-0 showing highest crossover frequency (**Fig. 6B**, **Table S1**, and **Table S7**). In contrast to the other centromeres, the Cvi-0 hybrid had the lowest crossover frequency, suggestive of a genetic or epigenetic feature in this interval that suppresses recombination (**Fig. 6B–6C** and **Table S1**). While the Cvi-0 *CEN178* array was substantially larger than Col-0, the Tanz-1 array was larger still and showed significantly higher crossover frequency than inbreds (**Fig. 6B-6C**, **Fig. S8**, **Table S1**, and **Table S7**). Five hybrids showed significant segregation distortion in favour of the Col-0 chromosome, compared to inbreds, whereas one hybrid, ANGE-B-2, showed distortion against Col-0 (**Fig. 6B**, **Table S1**, and **Table S7**). As for the other centromeres, variation in centromere-proximal crossover frequency was not correlated with the incidence of segregation distortion, the size of the *CEN178* arrays, or the epigenetic landscape (**Fig. 6B-6C**, **Fig. S2, S8**, and **Table S1**).

### Centromere-proximal crossover frequency and segregation distortion compared across all five chromosomes

In total, we compared sixty hybrid combinations using twelve accessions per centromere, to inbred Col-0 FTL strains, for both crossover frequency and segregation distortion (**Figs. 2-6** and **Tables S3-S7**). Forty-nine of these combinations showed either significantly elevated (n=27), or decreased (n=22), centromere-proximal crossover frequency, compared to Col-0 inbreds, consistent with widespread modification of recombination by natural variation. The most consistently highest-recombining hybrids were the relict accessions Cvi-0 and IP-Alo-19, where four of five *CEN* intervals showed higher crossover than inbreds in each (**Table S1**). Whereas Ler-0 hybrids showed suppressed crossovers across four of five centromeres (**Table S1**), which may indicate the action of *trans-*modifying effects in these hybrids. Interestingly, despite being more genetically diverged, the relict accessions showed proportionally more high-recombination hybrids, and fewer with low recombination, than the Eurasian accessions (**Table S1**). *CEN3* and *CEN4* hybrids showed a higher frequency of hybrids with suppressed crossover events compared to the other centromeres; and *CEN4* in particular showed eleven of twelve hybrids with reduced crossovers compared to inbreds (**Table S1**), which we propose is due to the knob-inversion in Col-0 [50,51] (**Fig. S7**). Together, this indicates widespread modification of proximal crossover frequency, although with accession- and centromere-specific effects.

Compared to crossover frequency, we observed fewer (18 of 60) instances of hybrids showing segregation distortion, compared to inbreds (**Table S1**). There was no correlation between changes to crossover frequency and the incidence of segregation distortion (**Fig. S3** and **Table S1**), indicating these phenomena are uncoupled. We observed eighteen instances of the Col-0 FTL chromosomes over-transmitting, and five cases where they are under-transmitted, with a greater incidence on the longer chromosomes one and five (**Table S1**). Proportionally, there were more incidences of drive amongst the diverged relict hybrids, compared to the Eurasians, and the Tanz-1 accession showed distortion in favour of Col-0 across four of five centromeres. However, the strength of segregation distortion did not correlate with the size difference of hybrid centromeric *CEN178* arrays (**Fig. S3-S8**).

Finally, we investigated whether polymorphisms in CENH3 might contribute to variation in segregation distortion between hybrids (**Fig. S9**). We observed few amino acid polymorphisms between the Col-0 CENH3 sequence and those present in the accessions crossed to, with a G to A substitution in Elk-3 and IP-Cat-0, and a S to Y substitution in Cvi-0 (**Fig. S9**). In contrast, we observe a high number of variants, clustered in the N-terminal tail of CENH3, between *A. thaliana* and the sister species *A. lyrata* (**Fig. S9**). We conclude that genetic variation in CENH3 is not able to explain the variation we observe in centromere-proximal crossover frequency and segregation distortion.

## Discussion

We surveyed the effects of natural variation on meiotic crossover frequency and segregation distortion, using twelve Arabidopsis hybrids, across the five centromeres. We observed widespread modification of crossover frequency in the majority of hybrids, with approximately equal cases where recombination was higher or lower than observed in inbreds. When considering genetic variation within the crossover-measured intervals, including the size and structure of the *CEN178* satellite arrays, there was no correlation between sequence divergence and recombination frequency. Indeed, the highest recombining accessions, Cvi-0 and IP-Alo-19, are both diverged relict accessions compared to Eurasian Col-0, indicating that *cis* genetic divergence does not explain variation in recombination levels. These observations are also consistent with previous work, where Arabidopsis hybrids show elevated recombination, compared to inbreds [52–54]. However, the knob-inversion polymorphism within *CEN4* of Col-0 provides a likely example of *cis* suppression by structural polymorphism [50,51]. As Cvi-0 and IP-Alo-19 showed high recombination across four of five centromeres, and Ler-0 showed low recombination across four of five, these may represent accessions with *trans*-acting recombination-modifying loci that exert a global effect on centromere-proximal crossovers. Further quantitative trait loci mapping experiments using populations derived from these hybrids may reveal new genes that regulate levels of centromeric crossover frequency.

Although less frequent than centromere-proximal crossover modification, we also observed multiple instances of segregation distortion across the five centromeres, in a minority of hybrids, compared to inbreds. Meiotic drive of centromeres has been observed in female meiosis of mice, and monkeyflowers, where only one of the meiotic products survives to produce the egg cell [32,33,36–39,55–57], as is known to occur in Arabidopsis (**Fig. 7**) [58,59]. Asymmetries in the spindle, or cell division, during female meiosis can be exploited by variant centromeres to cause segregation distortion into the surviving meiocyte [32,33,36–39,56,57]. Further work will be required to study Arabidopsis female meiosis in the developing ovule, and whether developmental or cellular asymmetries exist that driving centromeres may respond to (**Fig. 7A**) [58,59]. As we observe examples of segregation distortion, uncoupled from changes to centromere-proximal crossover frequency, these phenomena do not appear to be directly connected.

**Figure 7.**
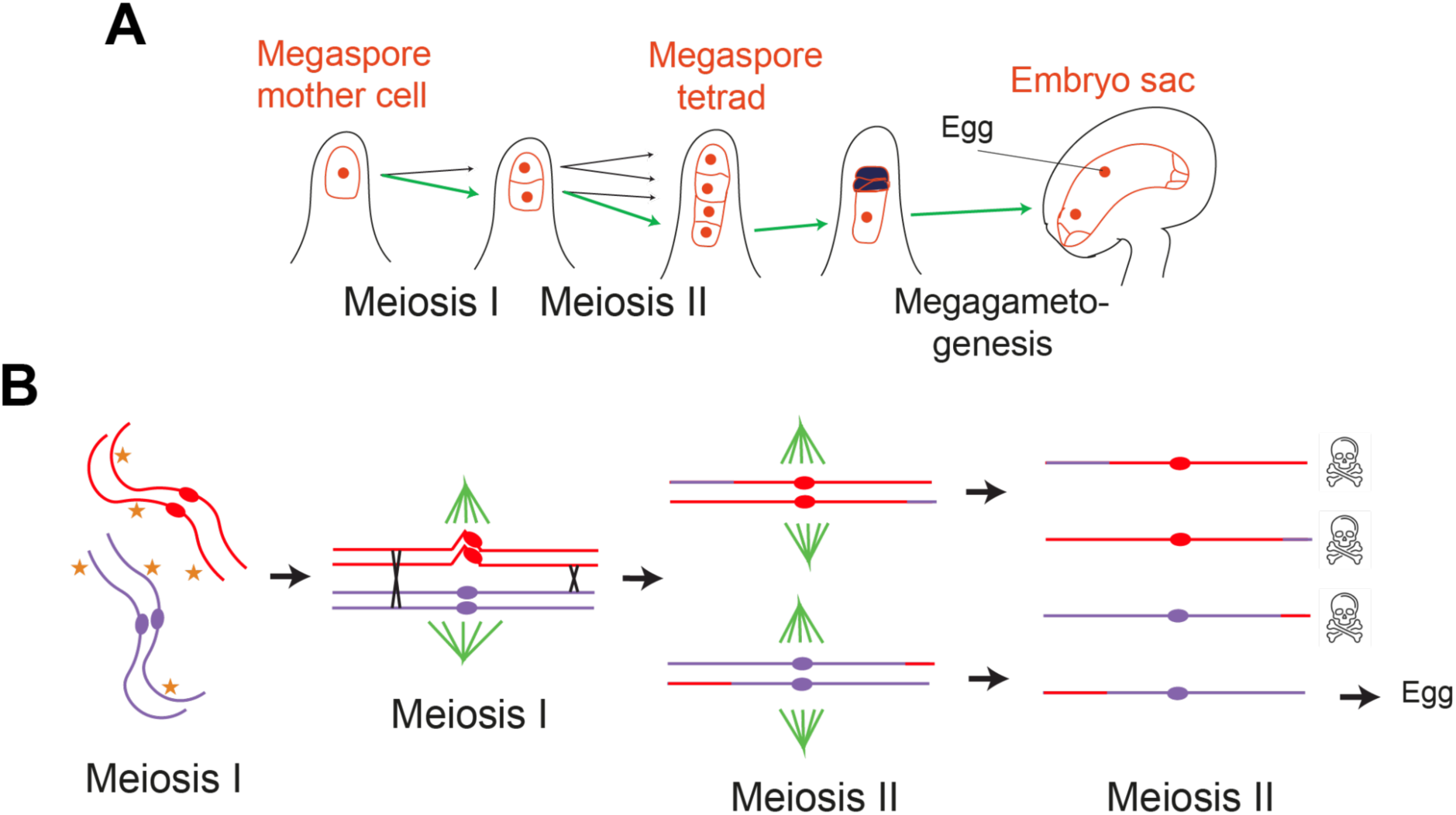
Model for how centromere structural polymorphism may cause segregation distortion in Arabidopsis female meiosis. **A.** Diagram representing the developing female ovule in Arabidopsis. A sub-epidermal cell becomes specified to enter meiosis, and undergoes two cell divisions to produce a tetrad of haploid spores. Three of the megaspores undergo programmed cell death, and the surviving cell tends to be the innermost spore [58,59]. The surviving megaspore then divides mitotically and differentiates into the female gametophyte, which contains the egg cell [58,59]. **B.** In early prophase-I in the sub-epidermal cell that is specified to enter meiosis, the homologous chromosomes (red and blue) replicate and are held together by cohesin and the meiotic axis. During prophase I, the replicated chromosomes experience DNA double strand breaks (DSBs, stars) that initiate meiotic recombination. As cell division progresses, the replicated homologs pair to form a four-strand bivalent structure, and a subset of the DSBs are repaired using a homologous chromatid to form crossovers (black crosses). At this stage we propose there is an asymmetry in the spindle microtubules, either in number or arrangement (green). Due to centromere structural heterozygosity, we propose that the synapsed homologs form a buckle where the sequence is hemizygous. Therefore, the relative positioning of the CENH3 chromatin (ovals) is offset between the homologs in the bivalent. In this case, this synapse-buckle and CENH3-offset cause a biased association of the purple centromeres with the lower spindle. The chromosomes then segregate through meiosis-I and meiosis-II, and because of the initial bias, the blue centromere has a higher chance of segregating into the surviving female spore that escapes programmed cell death and differentiates into the female gametophyte and egg cell.

We did not observe a clear correlation between genetic divergence, or the size of hybrid *CEN178* arrays, and the incidence of segregation distortion. However, as the region of CENH3 loading is of a consistent width in different accessions, with an enriched region of ∼1 Mb, irrespective of the size and structure of the underlying *CEN178* arrays, this may indicate that other features of centromere organisation are important for distorted segregation. For example, the relative placement of CENH3 regions on the recombining homologous chromosomes, rather than satellite array size, or the level of CENH3, may be more relevant for drive (**Fig. 7**). During meiosis, formation of DSBs are more frequent in the chromosome arms and are more likely to initiate interhomolog strand invasion and recombination earlier in prophase-I due to telomere-led synapsis [60,61]. As the Arabidopsis chromosome arms tend to be highly syntenic between accessions [21,40], they will synapse more effectively. We propose that the structurally polymorphic centromeres, which will form less DSBs, are likely to synapse in a delayed manner, causing a synapsis ‘buckle’ (**Fig. 7B**). Within these buckled centromeric regions, although a similar area is CENH3-occupied on both homologs, these regions may be positioned differently relative to the spindle microtubules, due to the effect of structural polymorphism on synapsis in the context of the paired bivalent structure (**Fig. 7B**). We speculate that such relative spatial offsets of CENH3 loading, and their differential interaction with proposed spindle asymmetries during female meiosis, may underlie the centromeric segregation distortion that we observe in Arabidopsis hybrids (**Fig. 7A-7B**). We also note that multiple instances of segregation distortion have been reported in Arabidopsis, suggesting such interactions could be widespread and result from interactions between *cis* and *trans*-acting factors [62–64]. Future studies that explore the cell biology of Arabidopsis meiosis, spindle formation, and how hybrid centromeres interact during pairing and recombination, may reveal the mechanisms at work.

## Materials and Methods

### Arabidopsis genetic stocks and growth conditions

All Col-0 FTL transgenic lines are TRAFFIC lines that were obtained from Prof. Scott Poethig (University of Pennsylvania) [44,46]. The ANGE-B-2 and ANGE-B-10 accessions were provided by Fabrice Roux (CNRS, Toulouse), IP-Alo-19, IP-Cat-0 and IP-Med-0 were provided by Carlos Alonso-Blanco (CNB-CSIC, Madrid), Elk-3, Rabacal-1 and Tanz-1 were provided by Angela Hancock (Max Planck Institute for Plant Breeding, Cologne), and Jm-0, Ler-0, Cvi-0 and Etna-2 were obtained from the Arabidopsis stock centre. All plants were grown in controlled environment chambers at 20°C, 60% humidity and a 16-hour photoperiod.

### Fluorescent seed scoring and calculation of crossover frequency and segregation distortion

Seed harvested from FTL hemizygous plants were cleaned to remove plant debris and images of a seed monolayer were captured using a Leica DFC310 FX dissecting microscope, using brightfield, or with ultraviolet filters. Images were analysed using an adapted CellProfiler pipeline (version 4.2.5) [65], which identifies seeds as primary objects and assesses fluorescence intensity for each object. Thresholds for fluorescence intensity were manually set for each seed image set, following inspection of the seed fluorescence intensity histograms. FTL crossover frequency (centiMorgan, cM) was calculated using the formula:

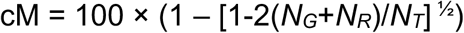

Where *N_G_* is a number of green-alone fluorescent seeds, *N_R_* is a number of red-alone fluorescent seeds, and *N_T_* is the total number of seeds counted, as described [52]. To test for significant differences in crossover frequency between the Col-0 inbred controls and each F_1_ hybrid, the *N_R_* and *N_G_* counts were summed across replicates for each genotype and combined with the remainder of total counts to construct 2×2 contingency tables and Chi-square tests performed. To test for significant differences in segregation distortion between the Col-0 inbred controls and each F_1_ hybrid, we compared the fluorescent and non-fluorescent seed counts to inbreds, using 2×2 contingency tables and Chi-square tests.

### Jm-0 and Elk-3 long-read DNA sequencing, genome assembly and annotation

A single Jm-0 individual was grown at 23°C under 16 hours of light and harvested 26 days after germination. 300 mg of tissue powder was used for high-molecular weight genomic DNA extraction. The methods for DNA extraction, PacBio HiFi library preparation, genome assembly using hifiasm (Version 0.16.1-r375; [48]), and quality assessment were extensively detailed in the previous release of the other ten HiFi- based assemblies used in this study [21].

For Elk-3 genomic DNA extraction associated with ONT sequencing, 3-week-old seedlings were grown on ½ MS media and 1% sucrose and kept in the dark for 48 hours prior to harvesting. Approximately 2 grams of tissue was used per extraction. Tissue was flash frozen and ground in liquid nitrogen, using a pestle and mortar for approximately 20 minutes. DNA was extracted using NucleoBond HMW DNA kit (MACHEREY-NAGEL), as per the manufacturer’s instructions. Library preparation followed the Nanopore SQK-LSK114 protocol and kit with the following modifications. Approximately 1.2-1.5 µg of size-selected DNA in a volume of 48 µl was used for library preparation. The first bead clean-up after nic-repair and end-prep was replaced by Short Read Eliminator (SRE) size selection (PacBio). Libraries were sequenced in a R10.4.1 Promethion flowcell. Base-calling and DNA methylation quantification was performed using dorardo (v0.8.3.), using model dna_r10.4.1_e8.2_400bps_sup@v5.0.0 and --modified-bases 5mC_5hmC.

For genome assembly, Elk-3 reads were filtered for length and accuracy using Filtlong (Version 0.2.0) (--min_mean_q 90, --min_length 50,000), and assembled using Hifiasm (Version 0.23.0-r691) using –ont parameters [48]. Chromosomes were generated as single contigs and extracted and oriented using Ragtag (Version v2.1.0) scaffolded to the Col-CEN assembly [28,66], and polished using medaka (version 2.0.1). For DNA methylation, reads were filtered for minimum length of 30 kb and aggregated using Modkit pileup (Version 0.4.1), using parameters (--ignore h --mod-thresholds C:0.85).

Copies of the centromeric satellite repeat were identified using TRASH, providing a template of the *CEN178* consensus sequence, as described [21,49]. To compare centromere structural and repeat polymorphism, we compared sequences using ReDotable for dot-plot analysis, as well as SyRI [67]. To map the location of synetic insertion sites for the FTL T-DNAs in the accession genome assemblies, we extracted 20 kb of sequence upstream and downstream of the insertion site from the Col-CEN assembly. These sequences were aligned to each of the accession genomes using LASTZ to identify the syntenic position in each genome. Protein sequences of the CENH3 coding region were extracted from the 13 studied *A. thaliana* accessions, and *A. lyrata* (MN47) [21], which encodes two CENH3 paralogues (Al1G11082 and Al1G59290) [68]. MAFFT (https://github.com/GSLBiotech/mafft) was used to align the protein sequences and snipit (https://github.com/aineniamh/snipit) was used for visualisation.

### Calculation of pairwise genetic diversity (π) in FTL regions

Genomes from each accession used in the crosses were aligned to the Col-CEN assembly [28], using Winnowmap (Version 2.03) with a k-mer size of 19 [69]. Sequence variant calling was performed using BCFtools (Version 1.21) [70]. Repetitive regions in the Col-CEN assembly were masked with RepeatMasker and TRASH, including centromeric satellites, transposable elements, organelle sequences, and rDNA [49]. Single nucleotide polymorphisms (SNPs) were filtered to exclude those overlapping repeat-masked regions, to retain strictly biallelic variants, and to limit missing data to no more than 20% across the examined accessions. Pairwise nucleotide diversity (π) relative to Col-CEN was calculated for each accession, within the positions of the FTL T-DNA markers on each chromosome.

### CENH3 ChIP-seq and analysis

Approximately 4 grams of 2-week-old seedlings were ground in liquid nitrogen. Nuclei were isolated in nuclei isolation buffer (1 M sucrose, 60 mM HEPES pH 8.0, 0.6% Triton X-100, 5 mM KCl, 5 mM MgCl2, 5 mM EDTA, 0.4 mM PMSF, 1 mM pepstatin-A, 1×protease inhibitor cocktail), and crosslinked in 1% formaldehyde at room temperature for 25 minutes. The crosslinking reaction was quenched with 125 mM glycine and incubated at room temperature for a further 25 minutes. The nuclei were purified from cellular debris via two rounds of filtration through one layer of Miracloth and centrifuged at 2,500g for 25 minutes at 4°C. The nuclei pellet was resuspended in EB2 buffer (0.25 M sucrose, 1% Triton X-100, 10 mM Tris-HCl pH 8.0, 10 mM MgCl2, 1 mM EDTA, 5 mM DTT, 0.1 mM 4 °C. PMSF, 1 mM pepstatin-A, 1×protease inhibitor cocktail) and centrifuged at 14,000g for 10 minutes at 4°C. The nuclei pellet was resuspended in lysis buffer (50 mM Tris-HCl pH 8.0, 1% SDS, 10 mM EDTA, 0.1 mM PMSF, 1 mM pepstatin-A), and chromatin was sonicated using a Covaris E220 evolution with the following settings: power=150V, bursts per cycle=200, duty factor=20%, time=60 seconds.

Sonicated chromatin was centrifuged at 14,000g and the supernatant was extracted and diluted with 1×volume of ChIP dilution buffer (1.1% Triton X-100, 20 mM Tris-HCl pH 8.0, 167 mM NaCl, 1.1 mM EDTA, 1mM pepstatin-A, 1×protease inhibitor cocktail). The chromatin was incubated overnight at 4°C with 50 µl Protein A magnetic beads (Dynabeads, Thermo Fisher) pre-bound with either 5 µl α-CENH3 raised to peptide acetyl-RTK HRV TRS QPR NQT DAC-amide (Eurogentec). The beads were collected on a magnetic rack and washed twice with low-salt wash buffer (150 mM NaCl, 0.1% SDS, 1% Triton X-100, 20 mM Tris-HCl pH 8.0, 2 mM EDTA, 0.4 mM PMSF, 1 mM pepstatin-A, 1×protease inhibitor cocktail) and twice with high-salt wash buffer (500 mM NaCl, 0.1% SDS, 1% Triton X-100, 20 mM Tris-HCl pH 8.0, 2 mM EDTA, 0.4 mM PMSF, 1 mM pepstatin-A, 1×protease inhibitor cocktail). Immunoprecipitated DNA– protein complexes were eluted from the beads (1% SDS, 0.1 M NaHCO3) at 65°C for 15 minutes. Samples were reverse crosslinked by incubating with 0.24 M NaCl at 65°C overnight. Proteins and RNA were digested with Proteinase K treatment, and RNase A, and DNA was purified with phenol:chloroform:isoamyl alcohol (25:24:1) extraction and ethanol precipitation. Library preparation followed the Tecan Ovation Ultralow System v2 library protocol. ChIP samples were PCR amplified for 12 cycles and sequenced with 150-bp paired-end reads on an Illumina instrument by Novogene.

### ChIP–seq data alignment and processing

Deduplicated paired-end CENH3 ChIP–seq Illumina reads (2 × 150 bp) from Col-0, Cvi-0, Ler-0 and Tanz-1 were processed with Cutadapt (v.1.18) to remove adaptor sequences and low-quality bases (Phred+33-scaled quality <20). For each accession, trimmed reads were aligned to the respective genome assembly using Bowtie2 (v.2.3.4.3), with the following settings: --very-sensitive --no-mixed --no-discordant -k 200 --maxins 1000. Read pairs with Bowtie2-assigned MAPQ of <10 were discarded using samtools (v.1.10). For retained read pairs that aligned to multiple locations, with varying alignment scores, the best alignment was selected. Alignments with more than two mismatches or consisting of only one read in a pair were discarded. For each dataset, bins per million mapped reads (BPM; equivalent to transcripts per million for RNA-seq data) coverage values were generated in bigWig and bedGraph formats with the bamCoverage tool from deepTools (v.3.5.0) and plotted using R.

### DNA methylation profiling and analysis

DNA base-calling, alignment and methylation calling was performed using dorardo (v0.8.3.) and model dna_r10.4.1_e8.2_400bps_sup@v5.0.0 and --modified-bases 5mC_5hmC. The reads were filtered for a minimum length of 30 kb, and aggregated using Modkit pileup (Version 0.4.1), using parameters (--ignore h --mod-thresholds C:0.85). The average methylation for each DNA sequence context was calculated over 10 kb windows, using R.

## Data availability

Long-read sequencing data and corresponding genome assemblies generated in this study for accessions Jm-0 and Elk-3 are available at the European Nucleotide Archive (ENA; https://www.ebi.ac.uk/ena/browser/home) under accession number PRJEB83900. Previously released long-read data and assemblies for the remaining then accessions are available at ENA under accession number PRJEB55353 [21].

## Acknowledgements

We acknowledge support from UKRI/BBSRC grants BB/Y009487/1 and BB/V003984/1, ERC grant 101142254 EvoPanCen, a UKRI/BBSRC DTP studentship to NG, and a Broodbank Fellowship to MN. We kindly thank Scott Poethig for providing the fluorescent TRAFFIC FTL reporter lines, and Fabrice Roux, Carlos Alonso-Blanco, and Angela Hancock for providing seed of Arabidopsis accessions.

---

**Figure S1.**
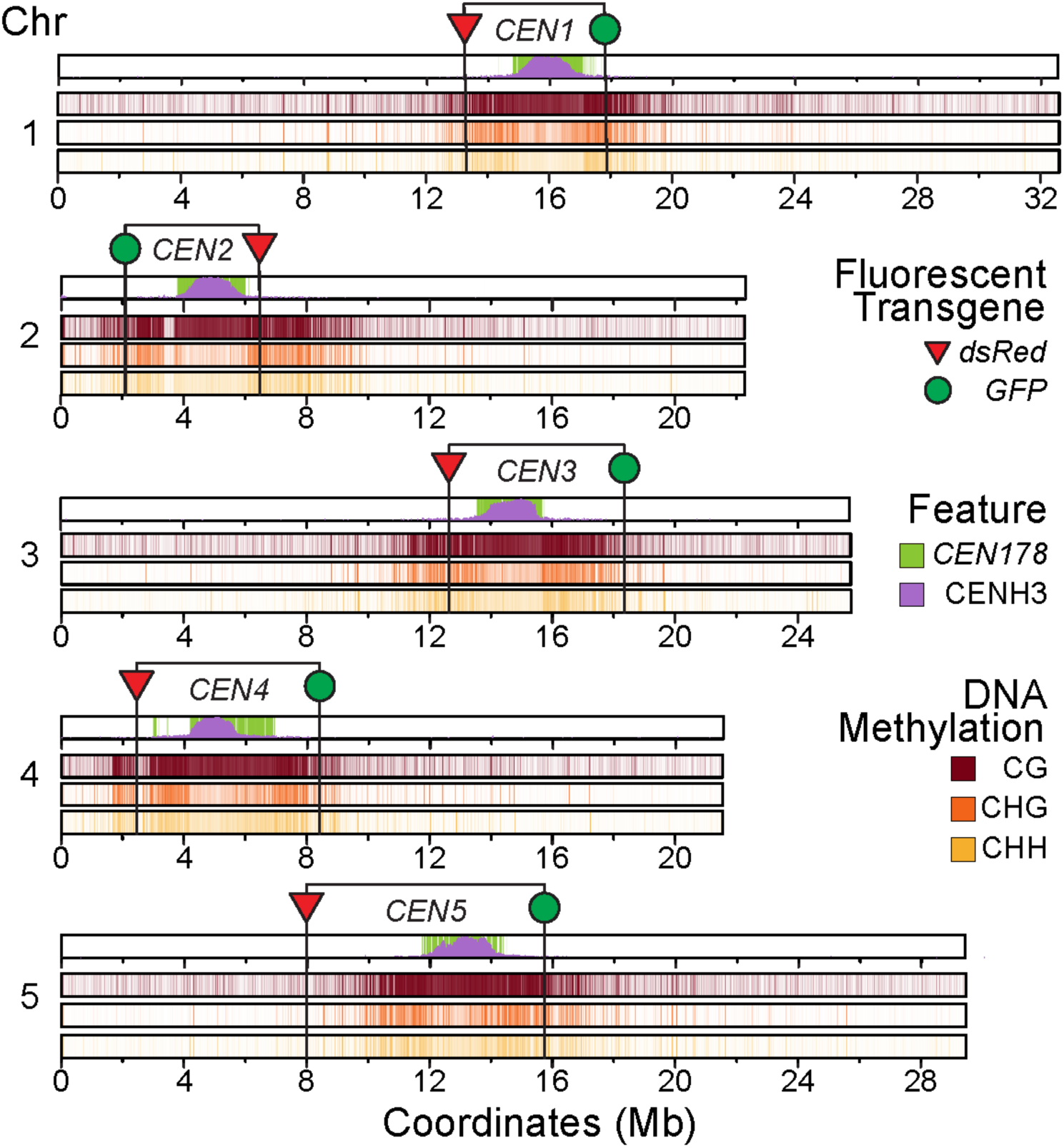
Location of *CEN* FTL intervals relative to the Col-0 genome and genetic and epigenetic features. For each chromosome in the Col-CEN assembly [28], we plotted the position of the FTL red-encoding (red triangles) and green-encoding (green circles) T-DNAs. Beneath we show density plots for (i) *CEN178* satellite repeats (green), CENH3 ChIP-seq enrichment (purple), and (ii) DNA methylation in CG (dark red), CHG (orange) and CHH (yellow) sequence contexts [21,28].

**Figure S2.**
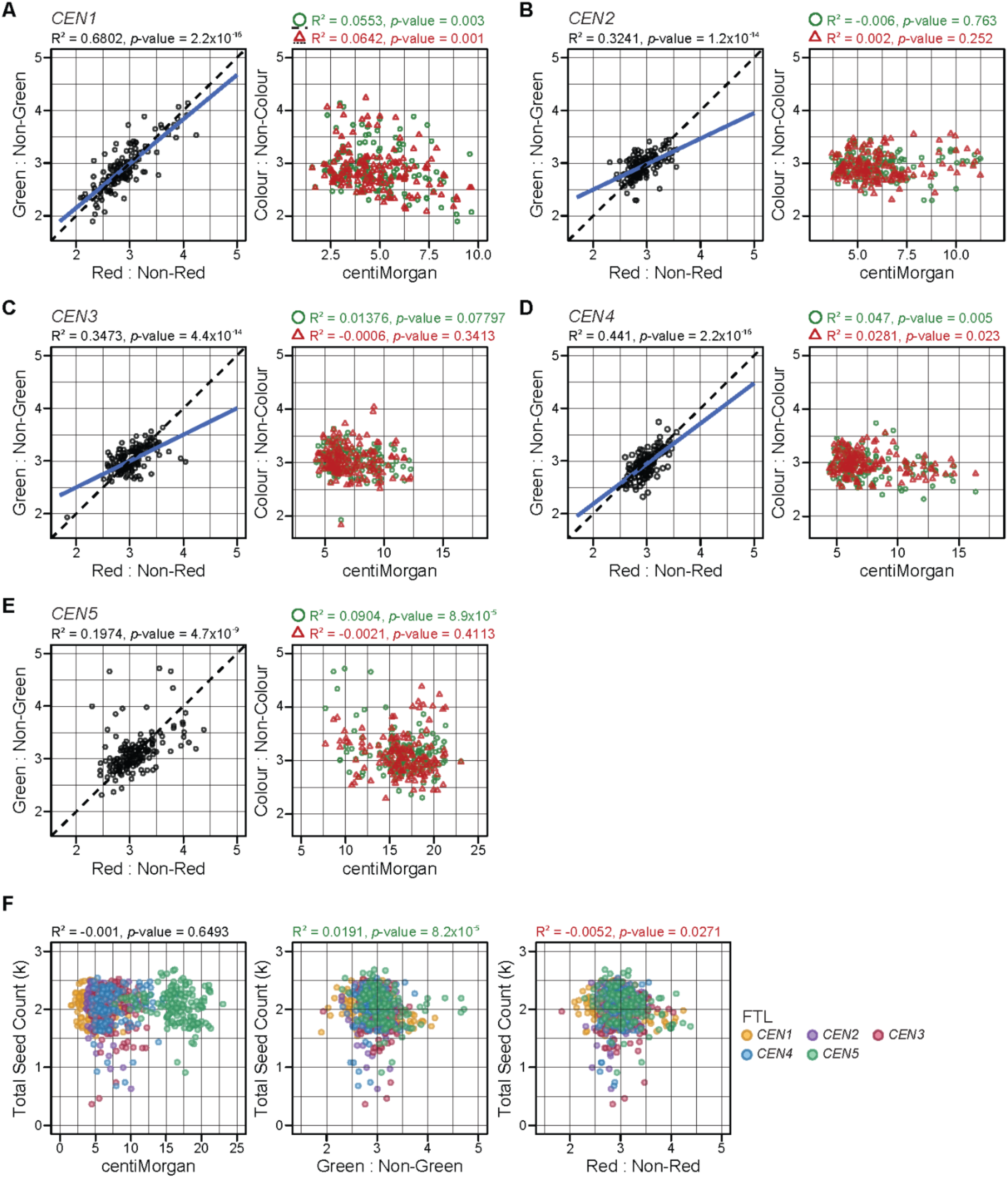
Correlation between FTL fluorescence ratios, crossover frequency and total seed counts. **A.** On the left, a scatter plot of green:non-green, and red:non-red *CEN1* seed fluorescence ratios, for every genotype and replicate. On the right, a scatter plot of colour:non-colour (red and green plotted separately) and *CEN1* crossover frequency (centiMorgan), for every genotype and replicate. Where the correlation is significant, a blue line is printed showing the relationship. Correlation values are printed above, together with the *P* value. **B.** As for A, but showing data for *CEN2*. **C.** As for A, but showing data for *CEN3*. **D.** As for A, but showing data for *CEN4*. **E.** As for A, but showing data for *CEN5*. **F.** As for A, but plotting total seed counts for each replicate against either centiMorgan, or green:non-green, or red-non-red fluorescence ratios, for each of the centromeres.

**Figure S3.**
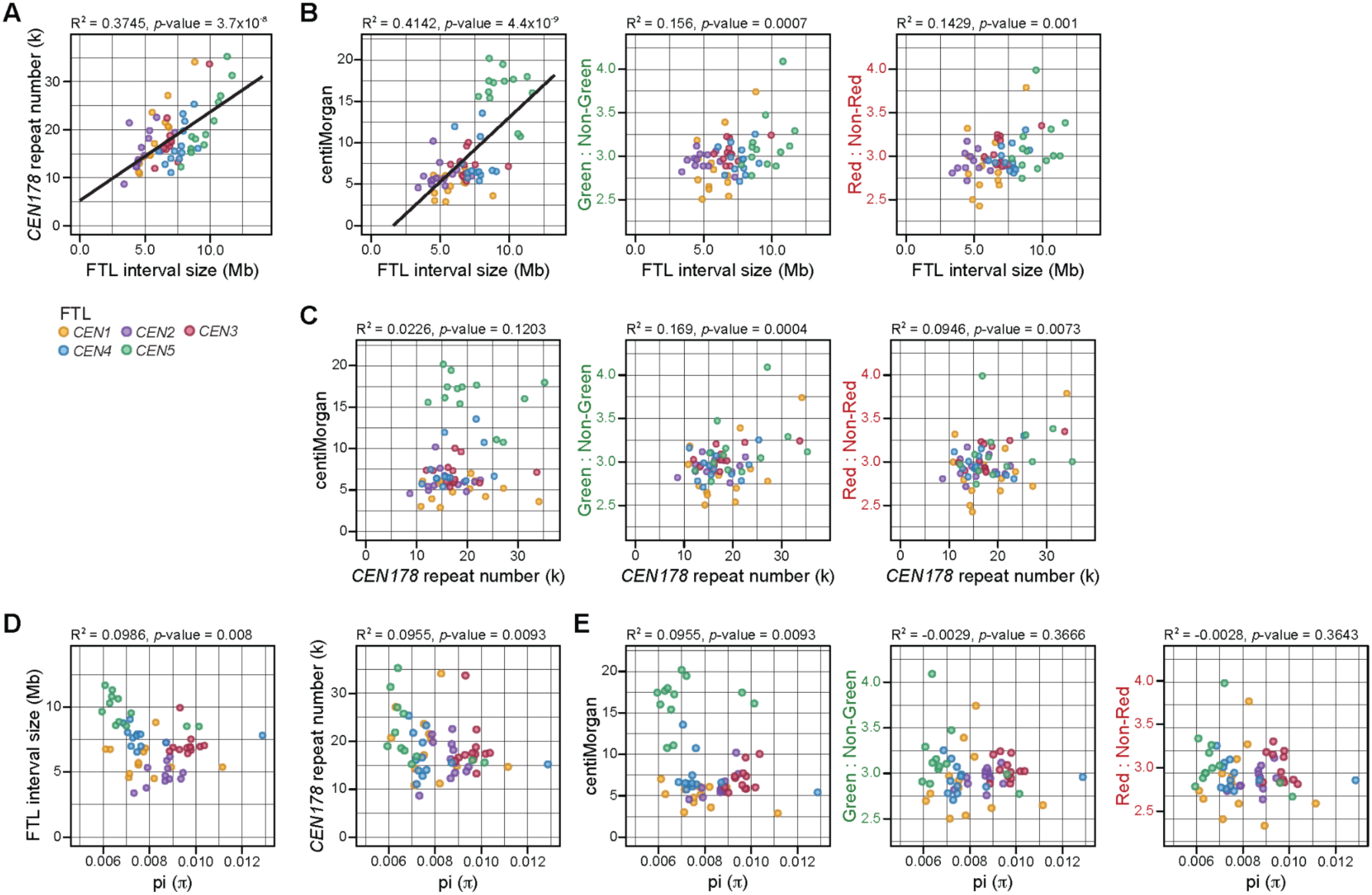
Correlation between FTL size, *CEN178* copies, crossover frequency, genetic diversity and fluorescence inheritance ratios. **A.** A scatter plot of *CEN178* repeat number annotated by TRASH, and the physical size of the FTL interval, in each accession. Data from the *CEN1*, *CEN2, CEN3*, *CEN4,* and *CEN5*, are coloured separately. Where the correlation is significant, a blue line is printed showing the relationship. Correlation values are printed above, together with the *P* value. **B.** As for A., but correlating FTL interval size with centiMorgans, or ratios of green:non-green, or red:non-red fluorescent seed counts. **C.** As for B, but comparing *CEN178* copy number. **D.** As for A, but pairwise SNP genetic diversity (π) in FTL regions, with FTL width, *CEN178* copy number, centiMorgans, or ratios of green:non-green, or red:non-red fluorescent seed counts.

**Figure S4.**
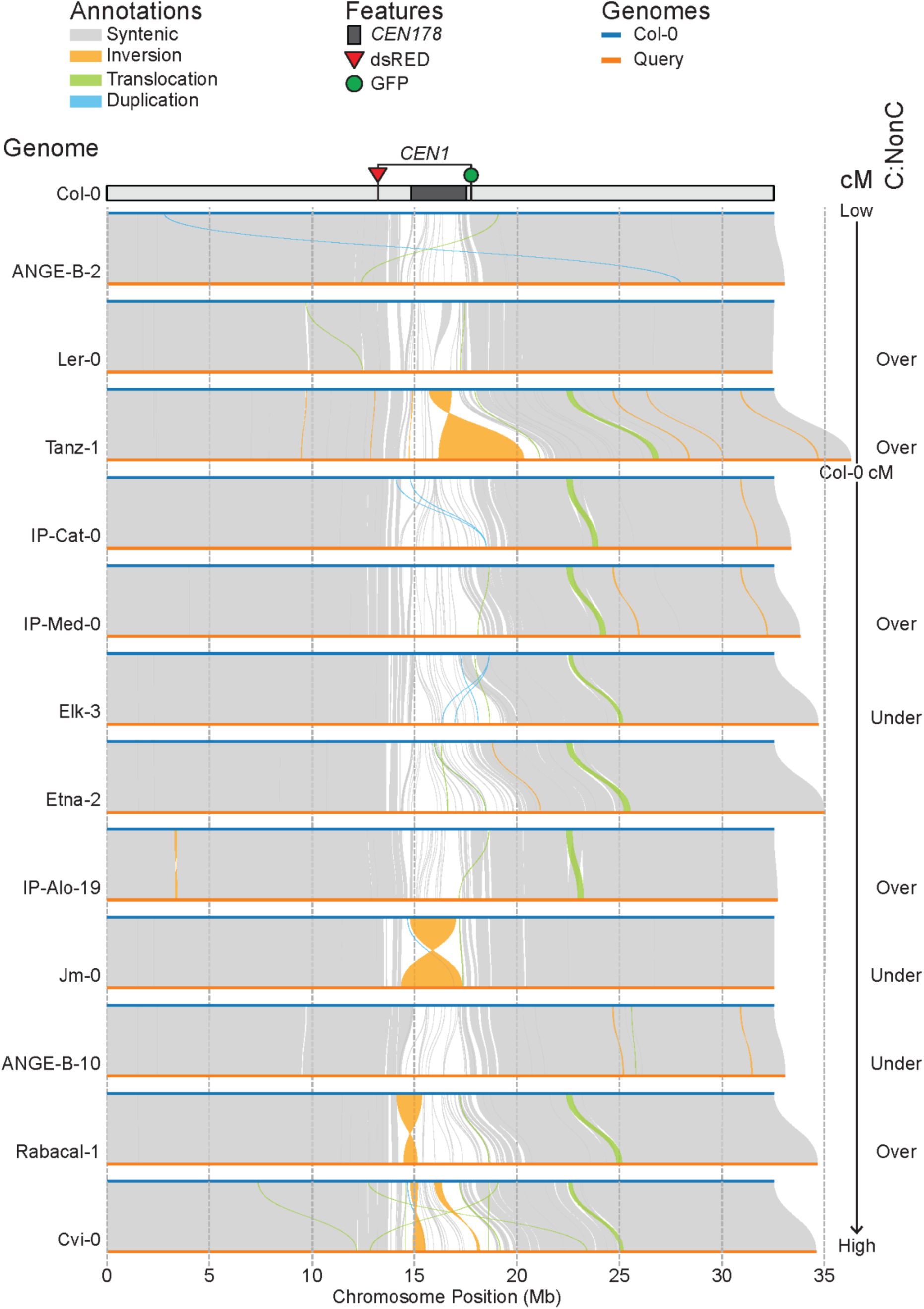
Patterns of genetic variation on chromosome 1 between Col-0 and the twelve hybrid accessions, ordered by *CEN1* crossover frequency. We used SyRI to map synteny between Col-CEN chromosome 1 (blue) and each of the accessions used to generate F_1_ FTL hybrids (orange) [67]. Regions detected by SyRI as syntenic (grey), inverted (orange), translocated (green), and duplicated (blue), are shaded between the chromosomes. The SyRI comparison plots are arranged from top to bottom in order of ascending *CEN1* crossover frequency in the cognate F_1_ hybrid. To the right, we also list whether there was over- or under-transmission of the *CEN1* FTL chromosome compared to inbreds. Segregation distortion is measured as colour:non-colour (C:NonC) ratios in the seed. At the top is a plot of Col-CEN chromosome 1 showing the positions of the FTL red-encoding (red triangle) and green-encoding (green triangle) T-DNAs, and the location of the centromeric *CEN178* satellite arrays (dark grey).

**Figure S5.**
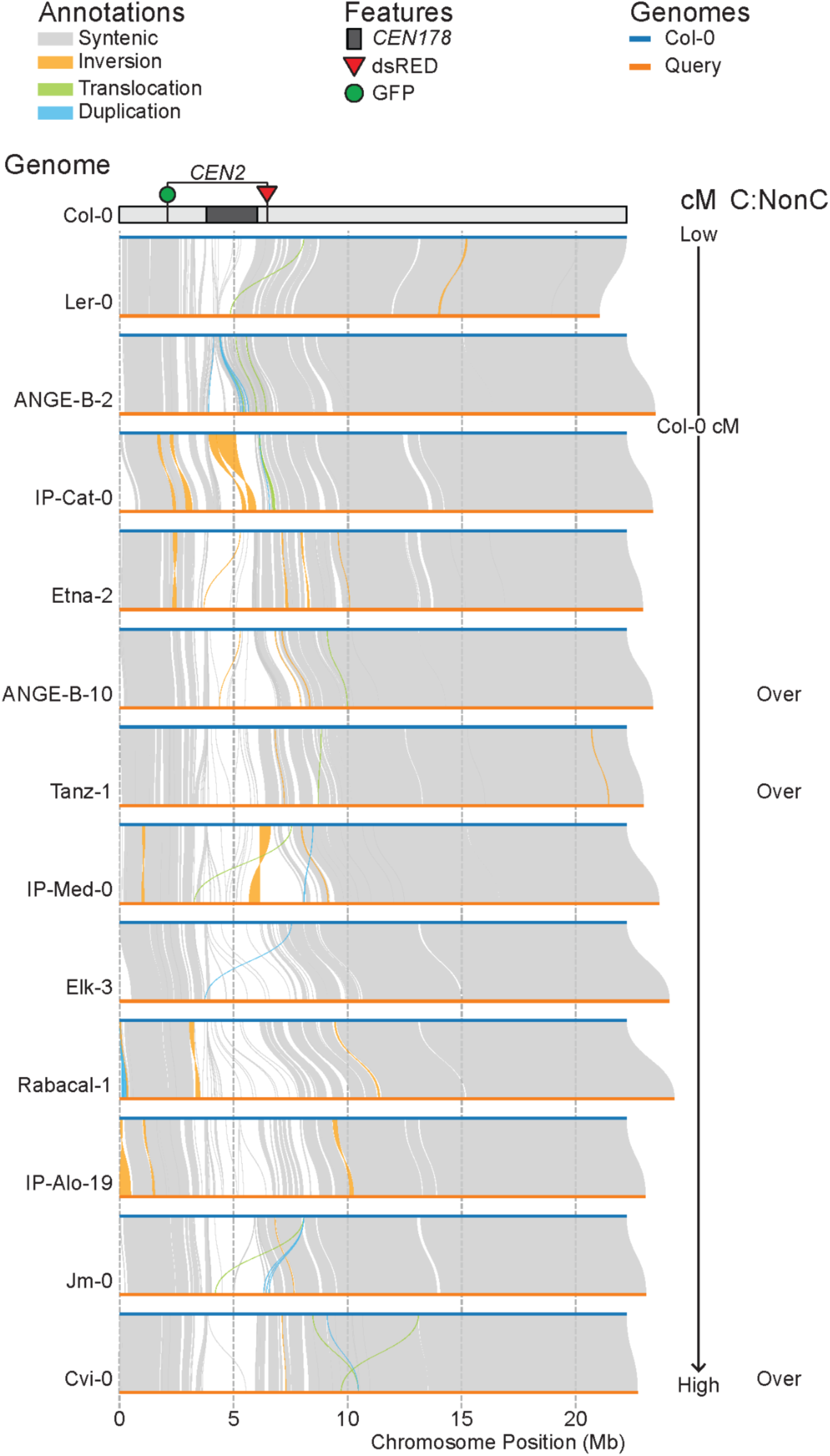
Patterns of genetic variation on chromosome 2 between Col-0 and the twelve hybrid accessions, ordered by *CEN2* crossover frequency. We used SyRI to map synteny between Col-CEN chromosome 2 (blue) and each of the accessions used to generate F_1_ FTL hybrids (orange) [67]. Regions detected by SyRI as syntenic (grey), inverted (orange), translocated (green), and duplicated (blue), are shaded between the chromosomes. The SyRI comparison plots are arranged from top to bottom in order of ascending *CEN2* crossover frequency in the cognate F_1_ hybrid. To the right, we also list whether there was over- or under-transmission of the *CEN2* FTL chromosome compared to inbreds. Segregation distortion is measured as colour:non-colour (C:NonC) ratios in the seed. At the top is a plot of Col-CEN chromosome 2 showing the positions of the FTL red-encoding (red triangle) and green-encoding (green triangle) T-DNAs, and the location of the centromeric *CEN178* satellite arrays (dark grey).

**Figure S6.**
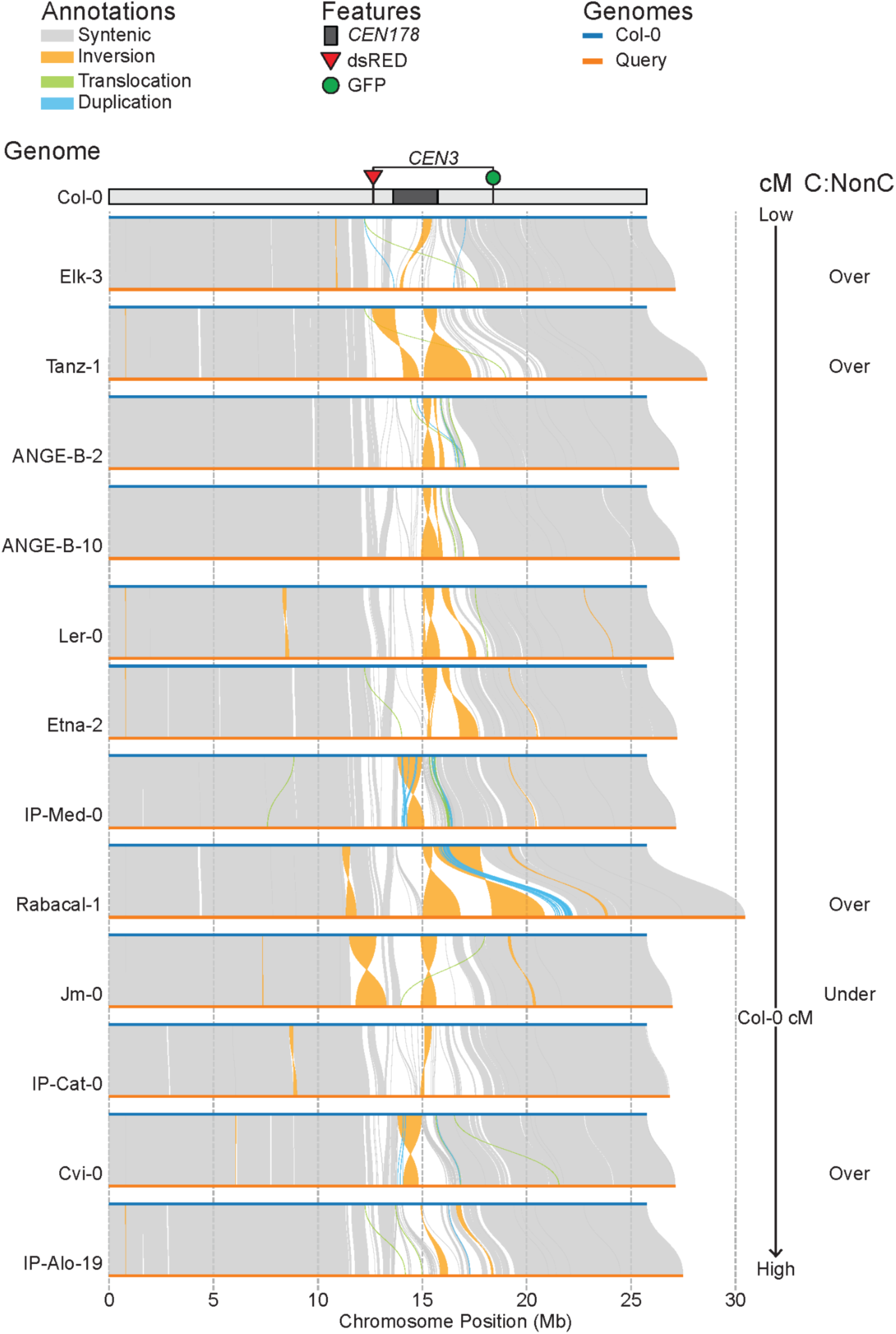
Patterns of genetic variation on chromosome 3 between Col-0 and the twelve hybrid accessions, ordered by *CEN3* crossover frequency. We used SyRI to map synteny between Col-CEN chromosome 3 (blue) and each of the accessions used to generate F_1_ FTL hybrids (orange) [67]. Regions detected by SyRI as syntenic (grey), inverted (orange), translocated (green), and duplicated (blue), are shaded between the chromosomes. The SyRI comparison plots are arranged from top to bottom in order of ascending *CEN3* crossover frequency in the cognate F_1_ hybrid. To the right, we also list whether there was over- or under-transmission of the *CEN3* FTL chromosome compared to inbreds. Segregation distortion is measured as colour:non-colour (C:NonC) ratios in the seed. At the top is a plot of Col-CEN chromosome 3 showing the positions of the FTL red-encoding (red triangle) and green-encoding (green triangle) T-DNAs, and the location of the centromeric *CEN178* satellite arrays (dark grey).

**Figure S7.**
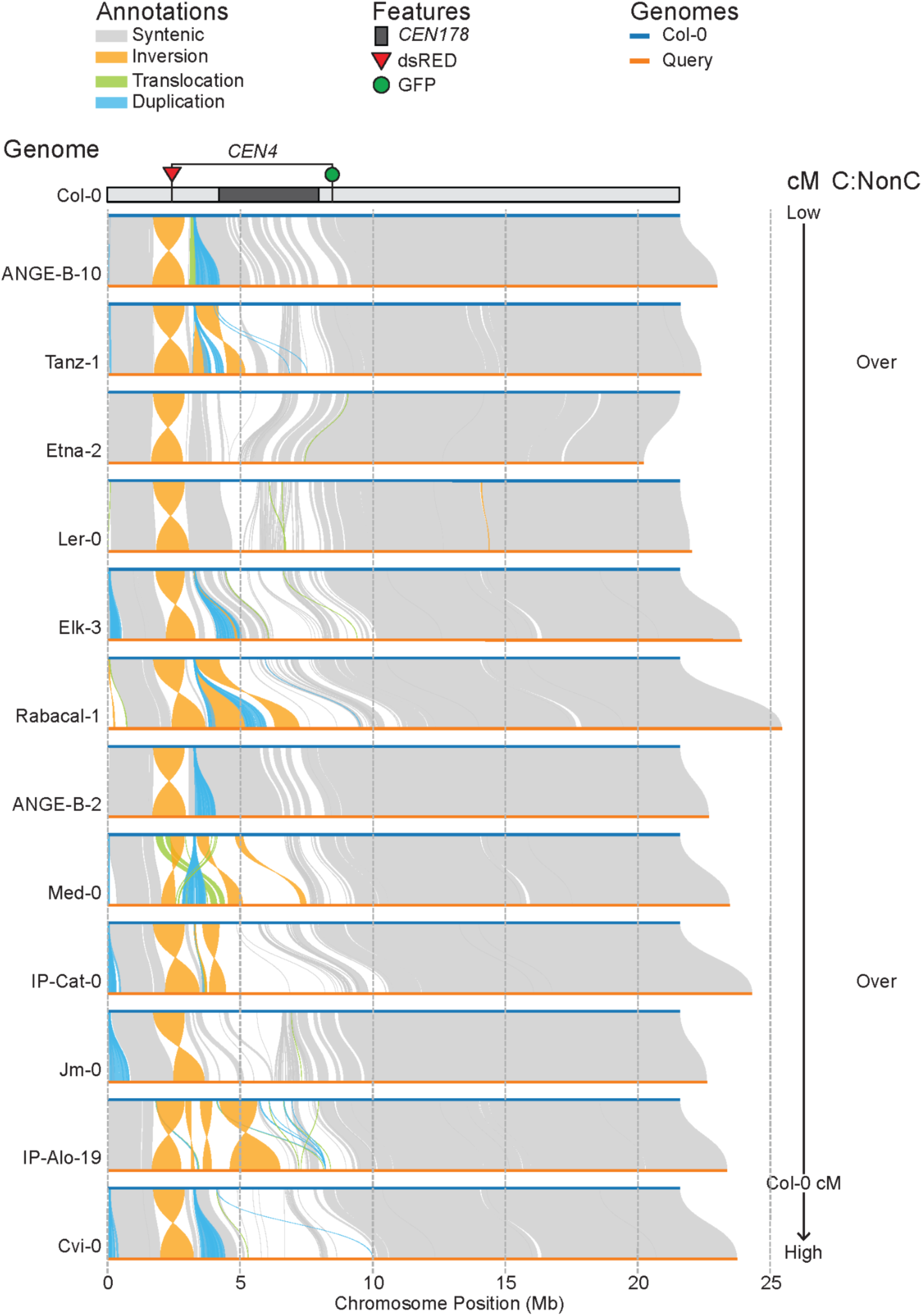
Patterns of genetic variation on chromosome 4 between Col-0 and the twelve hybrid accessions, ordered by *CEN4* crossover frequency. We used SyRI to map synteny between Col-CEN chromosome 4 (blue) and each of the accessions used to generate F_1_ FTL hybrids (orange) [67]. Regions detected by SyRI as syntenic (grey), inverted (orange), translocated (green), and duplicated (blue), are shaded between the chromosomes. The SyRI comparison plots are arranged from top to bottom in order of ascending *CEN3* crossover frequency in the cognate F_1_ hybrid. To the right, we also list whether there was over- or under-transmission of the *CEN4* FTL chromosome compared to inbreds. Segregation distortion is measured as colour:non-colour (C:NonC) ratios in the seed. At the top is a plot of Col-CEN chromosome 4 showing the positions of the FTL red-encoding (red triangle) and green-encoding (green triangle) T-DNAs, and the location of the centromeric *CEN178* satellite arrays (dark grey).

**Figure S8.**
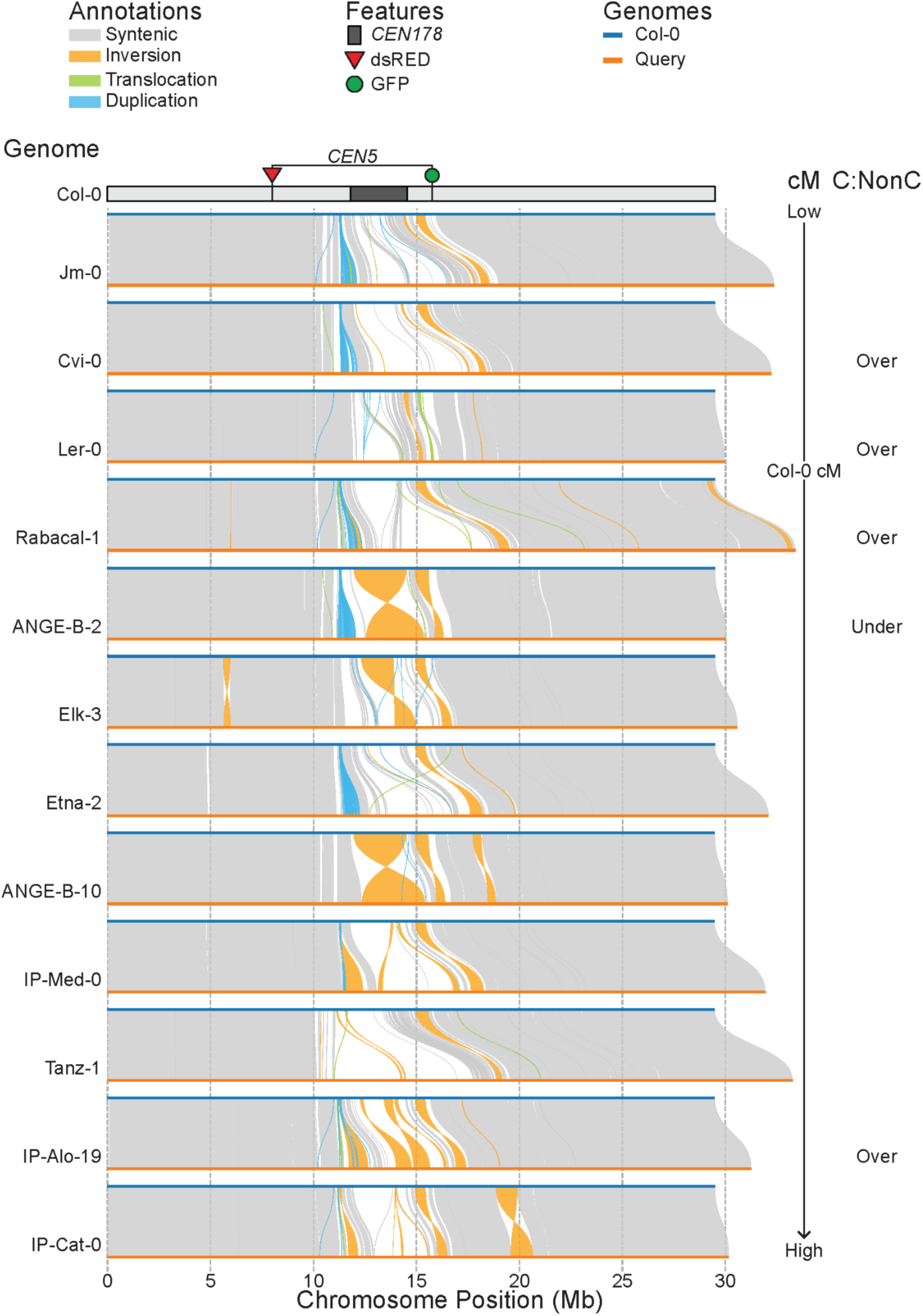
Patterns of genetic variation on chromosome 5 between Col-0 and the twelve hybrid accessions, ordered by *CEN5* crossover frequency. We used SyRI to map synteny between Col-CEN chromosome 5 (blue) and each of the accessions used to generate F_1_ FTL hybrids (orange) [67]. Regions detected by SyRI as syntenic (grey), inverted (orange), translocated (green), and duplicated (blue), are shaded between the chromosomes. The SyRI comparison plots are arranged from top to bottom in order of ascending *CEN5* crossover frequency in the cognate F_1_ hybrid. To the right, we also list whether there was over- or under-transmission of the *CEN5* FTL chromosome compared to inbreds. Segregation distortion is measured as colour:non-colour (C:NonC) ratios in the seed. At the top is a plot of Col-CEN chromosome 5 showing the positions of the FTL red-encoding (red triangle) and green-encoding (green triangle) T-DNAs, and the location of the centromeric *CEN178* satellite arrays (dark grey).

**Figure S9.**
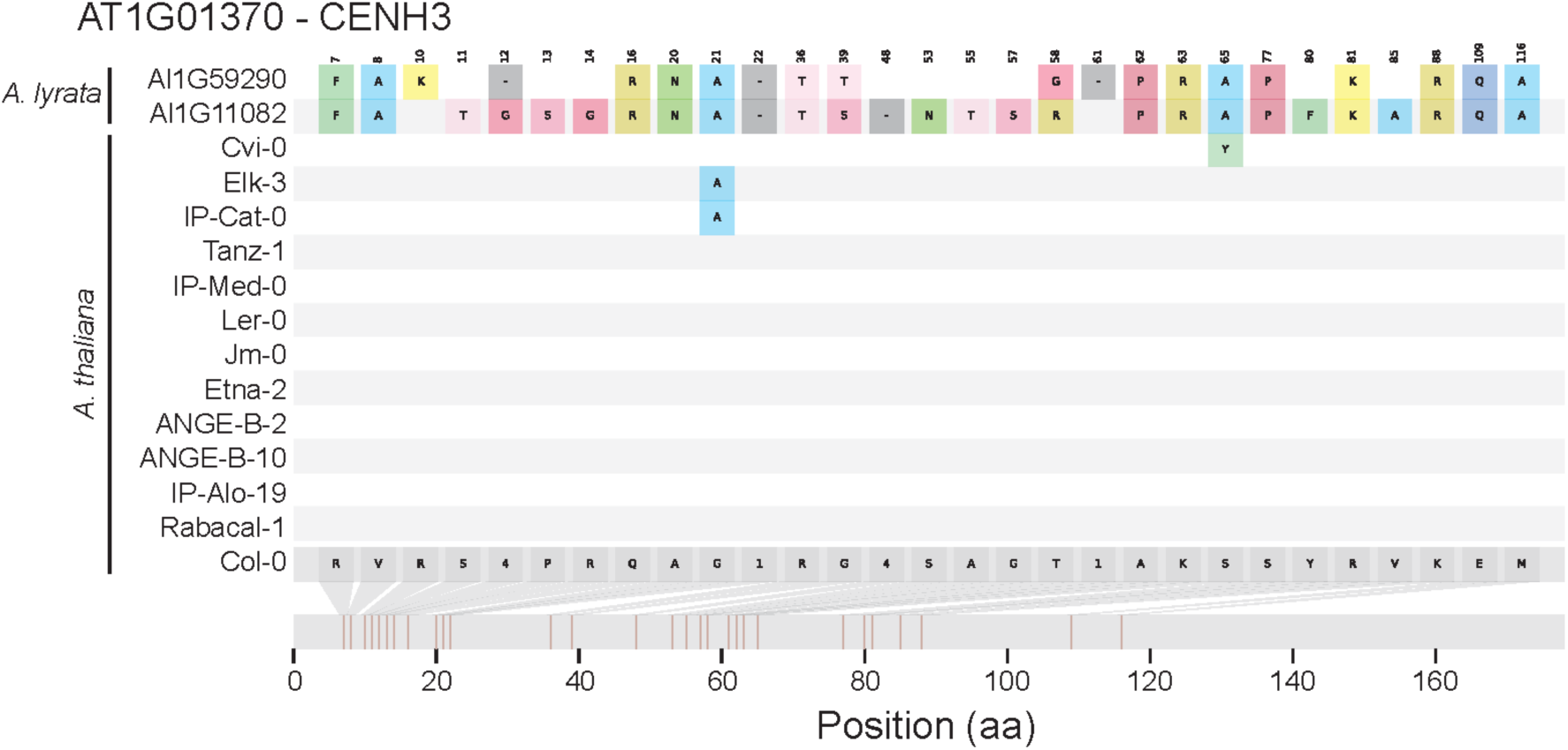
CENH3 amino acid variation present in the Arabidopsis accessions studied in comparison with *Arabidopsis lyrata*. Protein sequences of the CENH3 coding region were extracted from the 13 studied Arabidopsis accessions, and *A. lyrata* (MN47), which contains two CENH3 paralogues (Al1G11082 and Al1G59290) [68]. MAFFT (https://github.com/GSLBiotech/mafft) was used to align the protein sequences and snipit (https://github.com/aineniamh/snipit) was used for visualisation. Tick marks across the protein sequence represent polymorphisms compared to the reference Col-0 CENH3 sequence, and above is shown the reference amino acid. The position of the polymorphism and the amino acid change is shown at the top of the graph.

**Table S1. FTL interval size, annotation, crossover frequency, and segregation distortion for each centromere and hybrid.** For each FTL interval (*CEN1, CEN2*, *CEN3*, *CEN4* and *CEN5*), the accession name crossed to is listed, together with the associated genome assembly file name and which PCA genetic similarity group the accession belongs to (Eurasian, Iberian relict, or Non-Iberian relict) [21]. The row listing the Col-0 accession represents the inbred control line which the twelve hybrids are crossed to. FTL T-DNA flanking sequences from the Col-CEN assembly were mapped to each genome assembly and used to calculate FTL start and end coordinates, and the width of the interval in each accession. For each genotype the mean crossover frequency (centroMorgan, cM), and red:non-red-fluorescent (R:NR), and green:non-green-fluorescent (G:NG). The count data used for these calculations are provided in Tables S3-S7. Also listed are the number of *CEN178* satellite repeats annotated on this chromosome by TRASH [49], and the Jaccard similarity of the *CEN178* sequences present in each accession in each centromere with those in the Col-CEN assembly. Pairwise nucleotide diversity (π) relative to Col-0 was calculated for each accession using SNPs in the FTL T-DNA intervals on each chromosome.

**Table S2. *Arabidopsis thaliana* accessions analysed in this study.** Accession name is listed, together with chromosome arm SNP PCA genetic similarity group (Eurasian, Iberian relict or Non-Iberian relict), and the country, latitude and longitude of the collection location. We provide references for genome assemblies of these accessions used for analysis.

**Table S3. Fluorescent FTL seed count data for *CEN1*.** The table lists fluorescent seed count data from replicate hybrid F_1_ plants, where the accession listed was crossed to Col-0 *CEN1* lines. Counts of green-alone (*N_G_*), red-alone (*N_R_*), both red and green fluorescent and non-fluorescent seed, as well as the total number of seeds are listed, for each replicate. For each replicate, we also present crossover frequency (cM, centiMorgan), as well as the ratio of green to non-green fluorescent seed (G:NG), the ratio of red to non-red fluorescent seed (R:NR), and the ratio of green to red seed (G:R). The final column lists if genotypes were inbred or hybrid.

**Table S4. Fluorescent FTL seed count data for *CEN2*.** The table lists fluorescent seed count data from replicate hybrid F_1_ plants, where the accession listed was crossed to Col-0 *CEN2* lines. Counts of green-alone (*N_G_*), red-alone (*N_R_*), both red and green fluorescent and non-fluorescent seed, as well as the total number of seeds are listed, for each replicate. For each replicate, we also present crossover frequency (cM, centiMorgan), as well as the ratio of green to non-green fluorescent seed (G:NG), the ratio of red to non-red fluorescent seed (R:NR), and the ratio of green to red seed (G:R). The final column lists if genotypes were inbred or hybrid.

**Table S5. Fluorescent FTL seed count data for *CEN3*.** The table lists fluorescent seed count data from replicate hybrid F_1_ plants, where the accession listed was crossed to Col-0 *CEN3* lines. Counts of green-alone (*N_G_*), red-alone (*N_R_*), both red and green fluorescent and non-fluorescent seed, as well as the total number of seeds are listed, for each replicate. For each replicate, we also present crossover frequency (cM, centiMorgan), as well as the ratio of green to non-green fluorescent seed (G:NG), the ratio of red to non-red fluorescent seed (R:NR), and the ratio of green to red seed (G:R). The final column lists if genotypes were inbred or hybrid.

**Table S6. Fluorescent FTL seed count data for *CEN4*.** The table lists fluorescent seed count data from replicate hybrid F_1_ plants, where the accession listed was crossed to Col-0 *CEN4* lines. Counts of green-alone (*N_G_*), red-alone (*N_R_*), both red and green fluorescent and non-fluorescent seed, as well as the total number of seeds are listed, for each replicate. For each replicate, we also present crossover frequency (cM, centiMorgan), as well as the ratio of green to non-green fluorescent seed (G:NG), the ratio of red to non-red fluorescent seed (R:NR), and the ratio of green to red seed (G:R). The final column lists if genotypes were inbred or hybrid.

**Table S7. Fluorescent FTL seed count data for *CEN5*.** The table lists fluorescent seed count data from replicate hybrid F_1_ plants, where the accession listed was crossed to Col-0 *CEN5* lines. Counts of green-alone (*N_G_*), red-alone (*N_R_*), both red and green fluorescent and non-fluorescent seed, as well as the total number of seeds are listed, for each replicate. For each replicate, we also present crossover frequency (cM, centiMorgan), as well as the ratio of green to non-green fluorescent seed (G:NG), the ratio of red to non-red fluorescent seed (R:NR), and the ratio of green to red seed (G:R). The final column lists if genotypes were inbred or hybrid.

